# Deep learning models of cognitive processes constrained by human brain connectomes

**DOI:** 10.1101/2021.10.12.464145

**Authors:** Yu Zhang, Nicolas Farrugia, Pierre Bellec

**Author notes:** **Corresponding Authors:** Yu Zhang, Artificial Intelligence Research Institute, Zhejiang Lab, Zhongtai Street, Yuhang District, Hangzhou 311100, Zhejiang, China, Pierre Bellec, Département de Psychologie, Université de Montréal, 4565, Chemin Queen-Mary, Montréal (Québec) H3W 1W5.

## Abstract

Decoding cognitive processes from recordings of brain activity has been an active topic in neuroscience research for decades. Traditional decoding studies focused on pattern classification in specific regions of interest and averaging brain activity over many trials. Recently, brain decoding with graph neural networks has been shown to scale at fine temporal resolution and on the full brain, achieving state-of-the-art performance on the human connectome project benchmark. The reason behind this success is likely the strong inductive connectome prior that enables the integration of distributed patterns of brain activity. Yet, the nature of such inductive bias is still poorly understood. In this work, we investigate the impact of the inclusion of multiple path lengths (through high-order graph convolution), the homogeneity of brain parcels (graph nodes), and the type of interactions (graph edges). We evaluate the decoding models on a large population of 1200 participants, under 21 different experimental conditions, acquired from the Human Connectome Project database. Our findings reveal that the optimal choice for large-scale cognitive decoding is to propagate neural dynamics within empirical functional connectomes and integrate brain dynamics using high-order graph convolutions. In this setting, the model exhibits high decoding accuracy and robustness against adversarial attacks on the graph architecture, including randomization in functional connectomes and lesions in targeted brain regions and networks. The trained model relies on biologically meaningful features for the prediction of cognitive states and generates task-specific graph representations resembling task-evoked activation maps. These results demonstrate that a full-brain integrative model is critical for the large-scale brain decoding. Our study establishes principles of how to effectively leverage human connectome constraints in deep graph neural networks, providing new avenues to study the neural substrates of human cognition at scale.

## 1. Introduction

Modern imaging techniques, such as functional magnetic resonance imaging (fMRI), provide an opportunity to map the neural substrates of cognition in-vivo, and to decode cognitive processes non-invasively. Brain decoding has been an active topic since Haxby and colleagues first proposed the idea of using fMRI brain responses to predict the category of visual stimuli presented to a subject (Haxby et al., 2001). Nowadays, a variety of computational models have been proposed in the field, including multi-voxel pattern recognition (Haxby et al., 2014; Poldrack, 2011), linear regression models (Huth et al., 2012; Nishimoto et al., 2011), as well as nonlinear models such as deep artificial neural networks (Li and Fan, 2019; Wang et al., 2020). These decoding studies have been mainly focused on distinguishing the spatial patterns of brain activation within a small region of interest modulated by a few experimental tasks. Such brain decoders have gained many successes when tackling unimodal cognitive processes, in particular vision (Schrimpf et al., 2020), and focusing on specific brain regions, for example in the ventral visual stream network (Haan and Cowey, 2011). We recently proposed a graph neural network (GNN) model to decode high-order cognitive functions using distributed neural activity across large-scale brain networks (Zhang et al. 2021). This GNN model relied on a fixed human connectome as a static graph, embedded task-evoked brain activity as dynamic signal s on the graph, and integrated within-network context of spatiotemporal dynamics underlying cognitive processes through deep graph convolutions. We have shown that GNN can successfully decode a variety of cognitive tasks in a large population of healthy subjects, achieving high decoding performance on the Human Connectome Project (HCP) task benchmark (Zhang et al., 2021). However, it remains unclear how the choices of connectome priors and the interactions at different scales impact on large-scale cognition decoding.

Here, we investigate a form of graph convolution called ChebNet, which has the ability to propagate information over a relatively larger neighborhood on the graph and integrate neural activity in a multiscale manner, ranging from segregated brain activity from local areas (K=0), to information integration within the same brain circuit/network (K=1) as well as between multiple networks (K>1), and eventually towards the full brain. We propose a multi-domain decoding model using ChebNet graph convolutions and investigate how to implement human connectome constraints in the decoding pipeline, i.e. the implementation of multiscale functional integration and the construction of a proper graph architecture. The connectome constraints start with a brain parcellation, which divides the whole brain into hundreds of brain regions, and a brain graph that captures hierarchical and modular structures in brain organization. A variety of parcellation schemes have been proposed in the literature, see the review paper by (Eickhoff et al., 2018), which subdivide the entire cortex into non-overlapping regions based on connectivity profiles derived from diffusion tractography (Fan et al., 2016), functional organization (Yeo et al., 2011; Schaefer et al., 2018), or multimodal imaging features (Glasser et al., 2016). As the abstract representation of human connectome, a brain graph captures the network organization of the brain structure and function, by using anatomical and functional connectivity, in both healthy and diseased populations (Bassett and Bullmore, 2009; Bullmore and Sporns, 2009; Bullmore and Bassett, 2011). Brain atlas and connectivity are the two key components to define the nature of interactions in GNN, with the scale of functional interactions between areas specified by the path length of information propagation, i.e. K-order in ChebNet. Studies have shown that the edge-sparsified graphs achieved superior performance on graph learning benchmarks (Ye and Ji, 2021). Whether the edge-sparsified graphs outperform densely connected human connectomes in the field of large-scale cognitive decoding remains unknown.

In the current study, we evaluate the ChebNet decoding model on a large population of 1200 participants, under 21 different experimental conditions, acquired from the task-fMRI database from the Human Connectome Project (HCP). We explore the optimal choices of functional integration and graph architectures on this decoding benchmark, including the resolution or homogeneity of brain parcels (nodes), the type of interactions (edges), the inclusion of multiple path lengths on graphs (graph convolutions), and the sparsity of brain graphs. Moreover, we assess the robustness of brain decoding by introducing perturbations on the graph architecture, for instance network misspecifications due to random rewiring and node attacks. Lastly, we visualize the contributing salient neuroimaging features and the learned graph representations (i.e. activations of the last ChebNet layer) of the decoding model and compare them to the findings in the neuroscience literature.

## 2. Material and methods

### 2.1. fMRI Datasets and Preprocessing

We used the block-design task-fMRI dataset from the Human Connectome Project S1200 release (https://db.humanconnectome.org/data/projects/HCP_1200). The minimal preprocessed fMRI data in both NIFTI and CIFTI formats were selected. The preprocessing pipelines includes two steps (Glasser et al., 2013): 1) fMRIVolume pipeline generates “minimally preprocessed” 4D time-series (i.e. “.nii.gz” file) that includes gradient unwarping, motion correction, fieldmap-based EPI distortion correction, brain-boundary-based registration of EPI to structural T1-weighted scan, non-linear (FNIRT) registration into MNI152 space, and grand-mean intensity normalization. 2) fMRISurface pipeline projects fMRI data from the cortical gray matter ribbon onto the individual brain surface and then onto template surface meshes (i.e. “dtseries.nii” file), followed by surface-based smoothing using a geodesic Gaussian algorithm. Further details on fMRI data acquisition, task design and preprocessing can be found in (Barch et al., 2013; Glasser et al., 2013). The task fMRI database includes seven cognitive domains, which are emotion, gambling, language, motor, relational, social, and working memory. In total, there are 23 different experimental conditions. Considering the short event design nature of the gambling trials (1.5s for button press, 1s for feedback and 1s for ITI) which constrain the temporal resolution of our brain decoding pipeline (i.e. using a 10s time window), in the following experiments, we excluded the two gambling conditions and only reported results on the remaining 21 cognitive states. The detailed description of the tasks used in this study can be found in (Barch et al., 2013; Zhang et al., 2021) and is also listed in Table **1**.

**Table 1.** Scanning parameters and experimental designs of HCP task-fMRI dataset. The entire dataset includes in total 21 cognitive states and 14,895 functional runs across the six cognitive domains. By using a 10s-time window (i.e. 15 functional volumes at TR=0.72s), we cut long task trials into multiple data samples and resulted in 138,662 data samples of fMRI signals. These data samples constituted the whole dataset for model training and evaluation. Each functional run may contain multiple data samples of fMRI time-series depending on the number of task trials in the fMRI paradigm as well as the duration of each experimental condition. For instance, using a 10s-time window, we generated 10 data samples from one motor functional run (10 trials with one sample per trial) and 16 data samples from one working-memory functional run (8 trials with two samples per trial).

| Task Domains | #Subjects | #Runs | #Volumes<br>per run | #Effective<br>Sample<br>size | #Con<br>d | Min duration<br>per block (sec) |
| --- | --- | --- | --- | --- | --- | --- |
| Motor | 1083 | 2 | 284 | 21,670 | 5 | 12 |
| Language | 1051 | 2 | 316 | 36,164 | 2 | 10 |
| Emotion | 1047 | 2 | 176 | 10,470 | 2 | 18 |
| Social<br>cognition | 1051 | 2 | 274 | 21,020 | 2 | 23 |
| Working-memory | 1085 | 2 | 405 | 34,736 | 8 | 25 |
| Relational<br>processing | 1043 | 2 | 232 | 14,602 | 2 | 16 |

### 2.2. Decoding brain activity using graph convolution

As a representative model for brain organization, brain graph has been widely used in the neuroscience literature by associating nodes with brain regions and defining edges via anatomical or functional connections (Bullmore and Sporns, 2009). Graph Laplacian and graph convolution provides a generalized framework to analyze data defined on irregular domains, for instance social networks and brain networks. Thus, a non-linear embedding of brain activity can be learned to project the brain graph onto a low-dimensional representational space (Ortega et al., 2018), for instance mapping the gradients of brain organization (Margulies et al., 2016). We recently found that the convolutions on brain graph encoded the within-network interactions of neural dynamics in cognitive tasks (Zhang et al., 2021). In this study, we applied a generalized form of graph convolution by using high-order Chebyshev polynomials and explored the impact of the following factors: 1) high-order interactions to encode both within- and between-network interactions; 2) different graph architectures, ranging from local regions (spatial graph), to neural circuits (anatomical graph) and to functional networks (functional graph); 3) different brain atlases at various resolutions.

#### 2.2.1. Step 1: construction of brain graph

The decoding pipeline started with a weighted graph 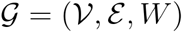, where 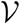 is a parcellation of cerebral cortex into *N* regions, 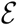 is a set of connections between each pair of brain regions, with its weights defined as *W_i,j_*. Many alternative approaches can be used to build such brain graph 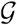, for instance using different brain parcellation schemes and constructing various types of brain connectomes. Here, we investigated multiple choices in both aspects. First, different parcellation schemes were used, including 1) functional subdivision of the cortical surface derived from resting-state functional networks (Yeo et al., 2011); 2) anatomical parcellation of both cortical and subcortical cortex derived from diffusion tractography (Fan et al., 2016); 3) a multimodal cortical parcellation bounded by sharp changes in cortical architecture, function, connectivity, and topography (Glasser et al., 2016); 4) a multi-scale atlas, consisting of brain parcels at different resolutions (Schaefer et al., 2018). Here, we used Glasser’s atlas as the default parcellation scheme, which achieved the highest performance on decoding 21 task states (see Results section). Second, we investigated various types of brain connectivity, i.e. edges between brain parcels, including 1) resting-state functional connectivity (RSFC); 2) anatomical connectivity (AC) derived from whole-brain tractography; 3) structural covariance (SC) of cortical thickness; 4) spatially adjacency (SP) in brain topology. Among which, the graph architecture derived from the anatomical and functional connectivity indicate biological or connectome constraints, while the spatial and structural graphs represent topological and morphological constraints respectively.

#### 2.2.2. Step 2: mapping of brain activity onto the graph

After the construction of brain graph (i.e. defining brain parcels and edges), for each functional run and each subject, we mapped the preprocessed task-fMRI data (e.g. “dtseries.nii” file for cortical parcellation, and “.nii.gz” file for volume parcellation) onto the set of brain parcels, resulting in a 2-dimensional time-series matrix. We created a task label with the same length as the fMRI time-series based on the experimental designs, e.g. task onsets and durations. Then, we used these task labels to extract a short series of fMRI responses from each functional run, by first splitting the entire run into multiple task blocks and then cutting the blocks into the chosen window sizes (discarding shorter time windows). To be noted that, each functional run usually contains multiple task blocks and each block is split into multiple bins of short time windows. As a result, we generated various numbers of fMRI time-series for each cognitive task, i.e. a short time-series with duration of *T* for each of *N* brain parcels, 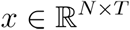, along with a unique task label for each time-series. In total, the entire dataset includes 14,895 functional runs across the six cognitive domains and over 1000 subjects for each domain, and results in 138,662 data samples of fMRI signals 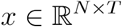 when using a 10s time window (i.e. 15 functional volumes at TR=0.72s), and 3,586,670 data samples when using a single-volume prediction.

#### 2.2.3. Step 3: spatiotemporal graph convolutions using ChebNet

Graph convolution relies on the graph Laplacian, which is a smooth operator characterizing the magnitude of signal changes between adjacent nodes. The normalized graph Laplacian is defined as:

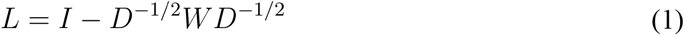

where *D* is a diagonal matrix of node degrees, *I* is the identity matrix, and *W* is the weight matrix which is specified by either multimodal brain connectivity (weighted graph) or the spatial adjacency in brain topology (binary graph). The eigendecomposition of Lapalcian matrix is defined as *L* = *U*Δ*U^T^*, where *U* = (*u*_0_,*u*_1_, ⋯ *u*_*N*–1_) is the matrix of Laplacian eigenvectors and is also called graph Fourier modes, and Δ = *diag*(*λ*_0_, *λ*_1_, ⋯ *λ*_*N*–1_ is a diagonal matrix of the corresponding eigenvalues, specifying the frequency of the graph modes. In other words, the eigenvalues quantify the smoothness of signal changes on the graph, while the eigenvectors indicate the patterns of signal distribution on the graph.

For a signal *x* defined on graph, i.e. assigning a feature vector to each brain region, the convolution between the graph signal 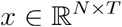 and a graph filter 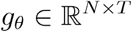 based on graph 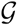, is defined as their element-wise Hadamard product in the spectral domain, i.e.:

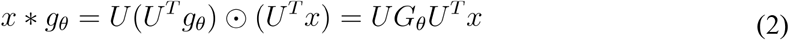

where *G_θ_* = *diag*(*U^T^g_θ_*) and *θ* indicate a parametric model for graph convolution *g_θ_*, *U* = (*u*_0_, *u*_1_, ⋯ *u*_*N*–1_ is the matrix of Laplacian eigenvectors and *U^T^x* is actually projecting the graph signal onto the full spectrum of graph modes. Equation (2) provided an easy way of calculating the graph convolution through a series of operations of matrix multiplication. To avoid calculating the spectral decomposition of the graph Laplacian, ChebNet convolution (Defferrard et al., 2016) uses a truncated expansion of the Chebychev polynomials, which are defined recursively by:

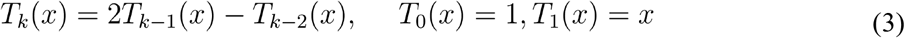

Consequently, the graph convolution is defined as:

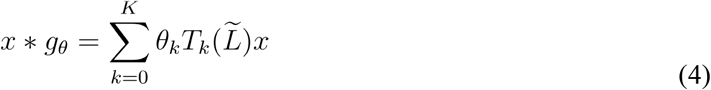

where 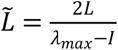 is a normalized version of graph Laplacian with *λ_max_* being the largest eigenvalue, *θ_k_* is the model parameter to be learned at each order of the Chebychev polynomials.

### 2.3. Brain-decoding pipeline

The proposed ChebNet decoding model (as shown in Figure **1**) consists of 6 graph convolutional layers with 32 graph filters at each layer, followed by a flatten layer and 2 fully connected layers (256, 64 units). The model takes in a short series of fMRI volumes as input, maps the fMRI data onto the predefined brain graph and results in a 2-dimensional time-series matrix 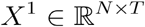, i.e. a short time-series with duration of *T* for each of *N* brain parcels. The first ChebNet layer learns various shapes of temporal convolution kernels by treating multiple time steps as input channels (*t*_0_, …, *t_T_*) and propagates such temporal dynamics within (K=1) and between (K>1) brain networks. As a result, a set of “brain activation” maps are generated and passed on to the second ChebNet layer for higher-order information integration, and so on. The learned graph representations at the last ChebNet layer (as shown in Fig. 9-S1c) are then imported into a 2-layer multilayer perceptron (MLP) for classification.

**Figure 1.**
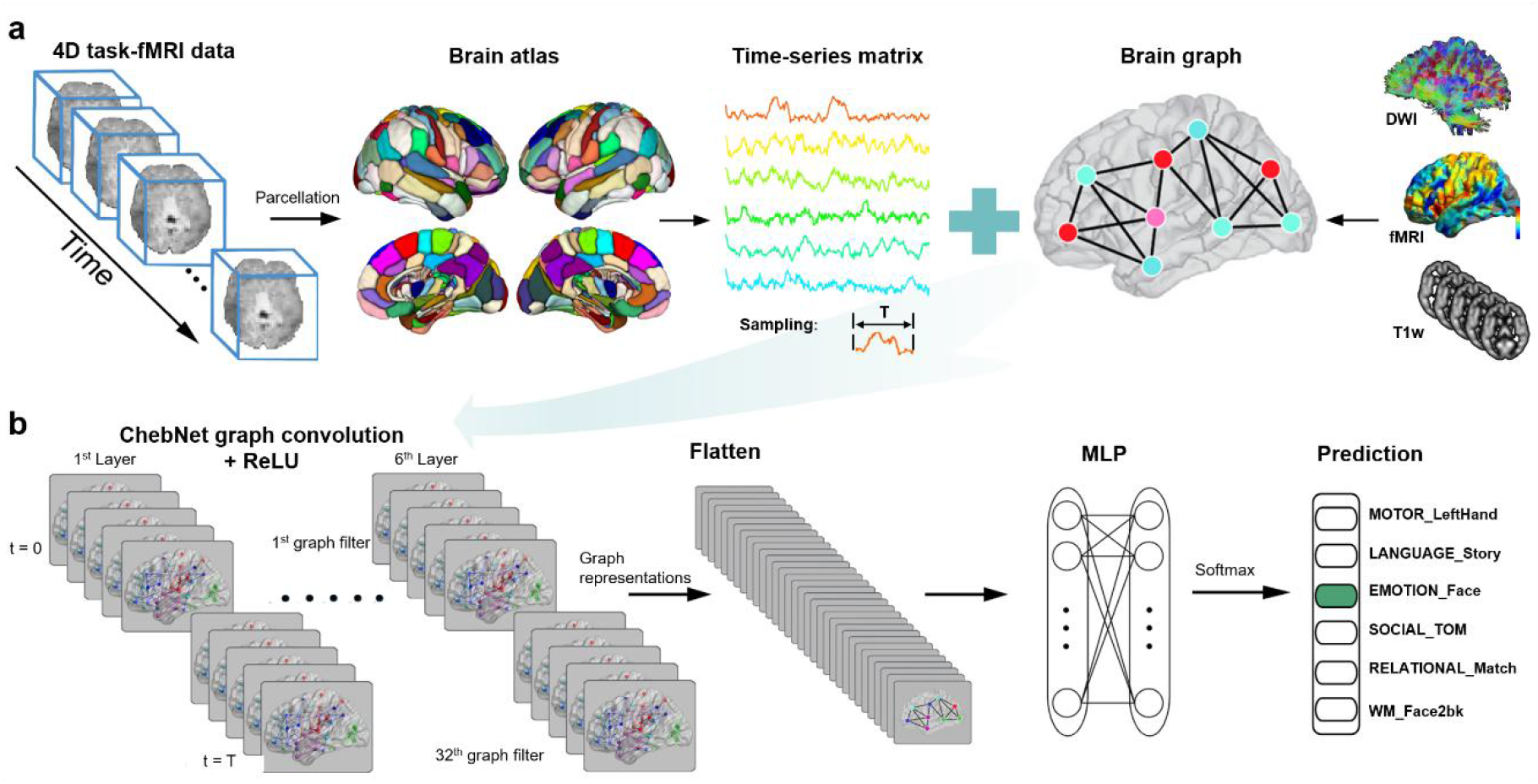
Pipeline of brain decoding using graph convolution network. We used a similar network architecture as proposed in our previous paper (Zhang et al., 2021) except for a more sophisticated form of graph convolution, namely ChebNet graph convolution, along with different brain graph architectures derived from various resolutions of brain atlas (nodes), and different types of brain connectivity (edges). The decoding model consists of six ChebNet graph convolutional layers with 32 graph filters at each layer, followed by a flatten layer and a two-layer Multilayer Perceptron (MLP, consisting of 256-64 units). Specifically, for a short series of fMRI volumes, the measured brain activity is first mapped onto a predefined brain atlas consisting of hundreds of brain regions (e.g. 246 regions from Brainnetome atlas (Fan et al., 2016)) and resulted in a 2D time-series matrix. Then, a brain graph indicating the edges between each pair of brain regions is constructed via either tractography of fiber projections using diffusion-weighted images (DWI), or functional correlation of low-frequency fluctuations in fMRI, or structural covariance of cortical thickness or gray matter density across a large population. Next, both the time-series matrix and brain graph are imported into the graph convolutional network (**b**). The model learns a new representation of task-evoked neural activity by stacking multiple graph convolutional layers, which takes into account both brain activity of each region (functional segregation) and the interactions within and between brain networks (functional integration). Finally, the learned graph representations are passed through a two-layer MLP and softmax function in order to predict the cognitive state associated with each input time window. The implementation of the ChebNet graph convolution was based on PyTorch 1.1.0, and was made publicly available in the following repository: https://github.com/zhangyu2ustc/gcn_tutorial_test.git.

The entire dataset includes in total 21 cognitive states and 14,895 functional runs across the six cognitive domains. By using a 10s-time window (i.e. 15 functional volumes at TR=0.72s), we cut long task trials into multiple data samples and resulted in 138,662 data samples of fMRI signals. These data samples constituted the whole dataset for model training and evaluation. The dataset was split into training (64%), validation (16%), test (20%) sets stratified by subject, which ensures that all fMRI data from the same subject was assigned to one of the three sets. Approximately, the training set includes fMRI data from 700 unique subjects (depending on data availability for different cognitive tasks ranging from 1043 to 1085 subjects in total), with 174 subjects for validation set and 218 subjects for test set. Specifically, the training set was used to train/update model parameters at each training epoch, the validation set was used to evaluate the model performance at the end of each training epoch, and the best model with the highest prediction score on the validation set was saved after 100 training epochs. The saved decoding model was then evaluated on the test set and reported the final decoding performance. We used Adam as the optimizer with an initial learning rate of 0.0001 on all cognitive domains. Additional l2 regularization of 0.0005 on weights was used to control model overfitting and the noise effect of fMRI signals. Dropout of 0.5 was additionally applied to the neurons in the last two fully connected layers. The implementation of the ChebNet graph convolution was based on PyTorch 1.1.0, and was made publicly available in the following repository: https://github.com/zhangyu2ustc/gcn_tutorial_test.git.

### 2.4. Effects of *K*-order in ChebNet

As stated in equation (4), the graph convolution can be rewritten as follows at different *K*-orders:

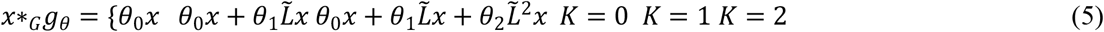

where 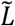 is a normalized version of graph Laplacian and {*θ_k_*}_*k*=1,2,..*K*_ are model parameters to be learned. Specifically, ChebNet graph convolution with K=0 indicates a global scaling factor on the input signal by treating each node independently; graph convolution with K=1 indicates information integration between the direct neighbors on graphs, i.e. integrating neural activity within the same brain network; graph convolution with K=2 indicates large-scale functional integration within a two-step neighborhood on graphs, i.e. integrating the context of neural activity from local areas, within- and between brain networks. Thus, the choice of K-order controls the scale of the information integration on graphs. Generally speaking, when K>1, the graph convolution integrates information flow within a *K*-step neighborhood by propagating graph signals not only within the same network but also among inter-connected networks. A simulation experiment shown in Figure 2-S1 illustrates that the propagation rate at all brain graphs converges to a similar level after *K*>10. For cognitive decoding, we explored different choices of *K*-order in ChebNet spanning over the list of [0,1,2,5,10] and found a significant boost in decoding performance by using high-order graph convolutions (*K*>1) instead of integrating within the network (*K*=1) especially on high-order cognition.

### 2.5. Definition of edges: brain connectivity

Graph architecture is another factor that impacts the propagation of information flow in the brain. We explored different types of edges (or brain connectivity) for brain decoding, including 1) local architecture indicating spatially adjacency in the cortical surface (spatial-graph); 2) community architecture representing neural circuits derived from whole-brain tractography (diffusion-graph); 3) hierarchical network architecture indicating functional correlation of low-frequency fluctuations in BOLD signals (functional-graph); 4) morphological networks derived from structural covariance of cortical thickness across a large population of subjects. These brain graphs were all calculated based on Glasser’s multimodal parcellation (Glasser et al., 2016). Specifically, for each pair of brain parcels in the atlas, the spatial-graph was evaluated by calculating the spatial adjacency matrix with each element indicating whether two brain parcels shared one or more triangles in the cortical surface by using the “wb_commond -cifti-label-adjacency” command from Connectome Workbench software. The diffusion-graph was generated from the whole-cortex probabilistic diffusion tractography based on HCP diffusion-weighted MRI data collected from 1065 subjects, with each element in the connectivity matrix indicating the average proportion of fiber tracts (streamlines) between the seed and target parcels (Rosen and Halgren, 2021). The functional-graph was calculated based on minimal preprocessed resting-state fMRI data from 1080 HCP subjects, by using the ‘tangent’ method for group-wise connectivity estimation with the ConnectivityMeasure method in nilearn (Varoquaux et al., 2010), and then averaged the connectivity matrices across all subjects. The structural-graph was generated by calculating structural covariance of cortical thickness based on the HCP structural MRI database across 1096 subjects. To be noted that, for spatial and structural graphs, only one brain graph was generated from the entire group. For the anatomical and functional graphs, we used the group averaged connectivity matrix and fixed the graph architecture for all subjects. Moreover, considering that the sparsification of graph is the key to superior performance on many graph learning benchmarks (Ye and Ji, 2021), a k-nearest-neighbour (k-NN) graph was built for each brain connectome by only connecting each brain region to its neighbours with the highest connectivity strength, resulting in an edge-sparsified brain graph. We have also explored how the sparsity of brain graphs impact brain decoding and compared them with the original densely-connected brain connectomes.

### 2.6. Definition of nodes: brain atlas

The parcellation scheme controls the scale of the graph. A variety of parcellation schemes have been proposed in the literature by using different imaging modalities and features (see the review paper by (Eickhoff et al., 2018)). We then investigated different parcellation schemes that have been widely used in the literature, with variable resolutions and connectivity information (e.g. functional, anatomical or multimodal connectivity information). For functional parcellation, considering the high correspondence between resting-state functional networks and patterns of task-evoked brain responses across multiple subjects, sessions, and cognitive tasks (Gordon et al., 2017; Gratton et al., 2018), we evaluated functional networks, for instance Yeo’s 7 and 17 resting-state networks consisting of 50 and 112 spatially continuous parcels in the cerebral cortex (Yeo et al., 2011), and functional parcellation at multiple resolutions, for instance Schaefer’s multiresolution brain parcellation, consisting of 100, 200, 400 and 1000 cortical parcels respectively (Schaefer et al., 2018). For anatomical parcellation, we chose the Brainnetome atlas that delineates the differences in connectivity profiles derived from diffusion tractography and consists of 246 brain regions in the cortical and subcortical areas (Fan et al., 2016). For parcellation with multimodal information, we chose the Glasser’s atlas that consists of 360 areas in the cerebral cortex, bounded by sharp changes in cortical architecture, function, connectivity, and topography (Glasser et al., 2016). For each parcellation map, we evaluated the functional homogeneity by calculating the averaged pairwise Pearson correlations within each brain parcel. We further investigated the relationship between the parcel size and functional homogeneity as well as the decoding performance among all parcellation maps.

### 2.7. Randomized brain graphs: edge rewiring

The robustness of the decoding model was investigated by introducing randomizations on both graph architectures and brain parcels. In order to evaluate whether the decoding model was impact by small fluctuations on the graph architecture, we generated a series of randomized null models that involves rewiring of the functional-graph by swapping a proportion of connections such that local degree is preserved while the global graph architecture is randomized (Sporns, 2018). These randomized null models can preserve graph attributes from the original graph, including local node measures, spatial locations, and wiring cost, and has been widely used as generative null models of the empirical data in network neuroscience literature, for instance (Betzel and Bassett, 2017). We explored different ratios of random swapping spanning over the list of [0, 0.1, 0.2, 0.5], where ratio=0.5 indicates that half of edges in the functional-graph were rewired. The randomized graphs to some degree represent the inherent mismatch between individuals or task-specific brain organization at a low ratio (e.g. 0.1), and may simulate disconnections in brain networks due to brain disorders by using a high ratio (Hearne et al., 2021; Sporns, 2011; Suárez et al., 2021). These randomized graphs were then compared with the original functional-graph in terms of decoding accuracy at different *K*-orders.

### 2.8. Randomized brain graphs: node attack and network attack

In order to evaluate whether the decoding model was robust to random or targeted attacks, we manually “silenced” a small portion of nodes (i.e. set their values to zero) and evaluated the reduction in the prediction accuracy by using pre-trained models. This procedure has been commonly used to simulate lesions due to brain injury or neurological disorders (Alstott et al., 2009; Honey and Sporns, 2008). For random node attack, we removed brain responses from randomly chosen nodes, ranging from 10% to 80% of brain parcels and repeated the process for 100 times. The vulnerability of each brain region was evaluated by associating the lesion state (i.e. lesion or normal) with the decays in brain decoding. For targeted or network attacks, we removed nodes associated with one intrinsic network at a time, identified by Yeo’s 7 or 17 resting-state functional networks (Yeo et al., 2011). Only brain regions with at least 30% areas affected were considered as lesions by silencing their activities in the following analysis. The vulnerability of each brain network was measured by the decay of decoding accuracy after applying network lesions as compared to the original decoding model.

### 2.9. Saliency map of graph convolutions

The saliency map analysis aims to locate which discriminative features in the brain help to differentiate between different cognitive tasks. There are several ways to visualize a deep neural network, including visualizing layer activation (Springenberg et al., 2014) and heatmaps (Selvaraju et al., 2020). Here, we chose the first method due to its easy implementation and generalization to graph convolutions (Springenberg et al., 2014). The basic idea is that if the input from a brain parcel is relevant to the prediction output, a little variation on the input signal will cause high changes in the layer activation. This can be characterized by the gradient of the output given the input, with the positive gradients indicating that a small change to the input signals increases the output value. Specifically, for the graph signal *X^l^* of layer *l* and its gradient *R^l^*, the overwritten gradient ∇_*X^l^*_*R^l^* can be calculated as follows:

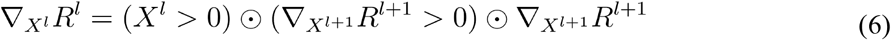

In order to generate the saliency map, we started with a pre-trained model and used the above chain rule to propagate the gradients from the output layer until reaching the input layer. This guided-backpropagation approach can provide a high-resolution saliency map with the same dimension as the input data. We further calculated a heatmap of saliency maps by taking the variance across the time steps for each parcel, considering that the variance of the saliency curve provides a simplified way to evaluate the contribution of task-evoked hemodynamic response. To make it comparable across subjects, the saliency value was additionally normalized to the range [0,1], with its highest value at 1 (a dominant effect for task prediction) and lowest at 0 (no contribution to task prediction).

## 3. Results

### 3.1. Decoding cognitive states with fine temporal resolution and high accuracy

We proposed a multi-domain decoding pipeline based on ChebNet graph convolution (Figure 1). The ChebNet decoding model was evaluated using the cognitive battery of HCP task-fMRI dataset acquired from 1200 healthy subjects. Using a 10-second window, the 21 cognitive states were identified with an average test accuracy of 93% (mean=93.43%, STD=0.44% by using 10-fold cross-validation with shuffle splits stratified by subject). The temporal resolution of the decoding model can go down to a single fMRI volume (720 ms of duration), with a prediction accuracy well above chance level (60%, chance level=4.8%). The accuracy of single-volume state annotation was highly improved (reaching 76%) after taking into account the delay effect of hemodynamic response function in BOLD signals, i.e. excluding fMRI volumes within 6s after the onset of each task trial from both training and test sets. We have also evaluated other baseline approaches including multi-class support vector machine classification (SVC) with linear and nonlinear kernels, random forest and a multilayer perceptron (MLP, consisting of two fully connected layers) on the same dataset. The results with a 10- second time window demonstrated that high-order ChebNet model outperformed all the other linear and nonlinear decoding models by a wide margin (see Table **2**). ChebNet also showed a significant improvement over GCN which relies on first-order graph convolution.

**Table 2:**
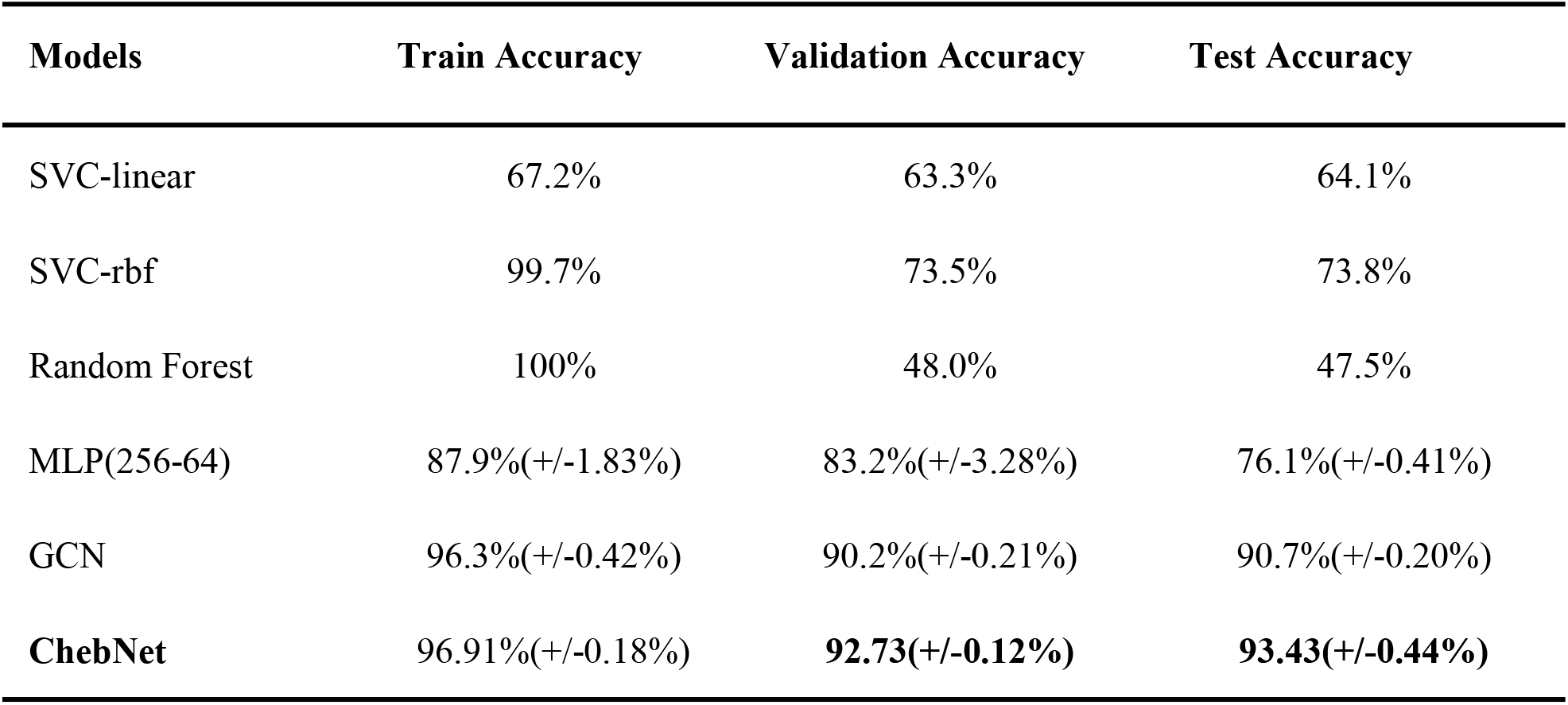
Comparison of cognitive decoding by using linear and nonlinear models. We reported the best performance for the baseline models after a grid search of the hyperparameters. For SVC approaches, we used the one-vs-rest (‘ovr’) decision function to handle multi-classes and reported the highest accuracy after the grid search for the hyper-parameter (C = [0.0001,0.001,0.1,1,10,100]). For Random Forest, we reported the highest accuracy after evaluating different settings of the classifier including depth of trees: [4,16,64,256,1024] and number of trees: [100,2000]. For MLP (multilayer perceptron), GCN (using first-order graph convolution, (Zhang et al., 2021)) and ChebNet (using 5-order graph convolution), we reported the mean and standard deviation of the decoding accuracies among 10-fold cross-validation with shuffle splits stratified by subject.

| Models | Train Accuracy | Validation Accuracy | Test Accuracy |
| --- | --- | --- | --- |
| SVC-linear | 67.2% | 63.3% | 64.1% |
| SVC-rbf | 99.7% | 73.5% | 73.8% |
| Random Forest | 100% | 48.0% | 47.5% |
| MLP(256-64) | 87.9%(+/-1.83%) | 83.2%(+/-3.28%) | 76.1%(+/-0.41%) |
| GCN | 96.3%(+/-0.42%) | 90.2%(+/-0.21%) | 90.7%(+/-0.20%) |
| **ChebNet** | **96.91%(+/-0.18%)** | **92.73(+/-0.12%)** | **93.43(+/-0.44%)** |

### 3.2. Low misclassification rate within and between cognitive domains

In order to clarify the nature of errors made by the ChebNet brain decoder, we examined the confusion matrix on the test set, which indicates the proportion of true and false predictions given a cognitive task state or domain. When using a 10-second time window, the confusion matrix showed a clear block diagonal architecture (see Figure **2a**), where the majority of experimental tasks were accurately identified for all task conditions (e.g. cross-domain misclassification rate<2%). Mistakes were found in tasks with similar cognitive processes, for instance, between relational processing and pattern matching conditions, as well as 0back vs 2back WM tasks. After summarizing the confusion matrix according to the six cognitive domains (Figure **2b**), each cognitive domain was identified with an accuracy greater than 95%. Among the six cognitive domains, the language tasks (story vs math) and motor tasks (left/right hand, left/right foot and tongue) were the most easily recognizable conditions by showing the highest precision and recall scores (average F1-score = 98% and 97%, respectively for two language conditions and five motor conditions). Even higher scores were achieved when decoding a smaller number of experimental conditions restricted to a specific cognitive domain (Figure 2-S1), i.e. using task-fMRI data exclusively from a single cognitive domain during model training and evaluation. For instance, the model achieved near-perfect decoding on other high-order cognitive functions, including working-memory (94.51%, classifying 8 conditions using 25s) and social cognition (96.58%, classifying 2 conditions using 23s), as opposed to 92.6% and 92.9% for the two domains when decoding on 30s of fMRI data (Li and Fan, 2019).

**Figure 2.**
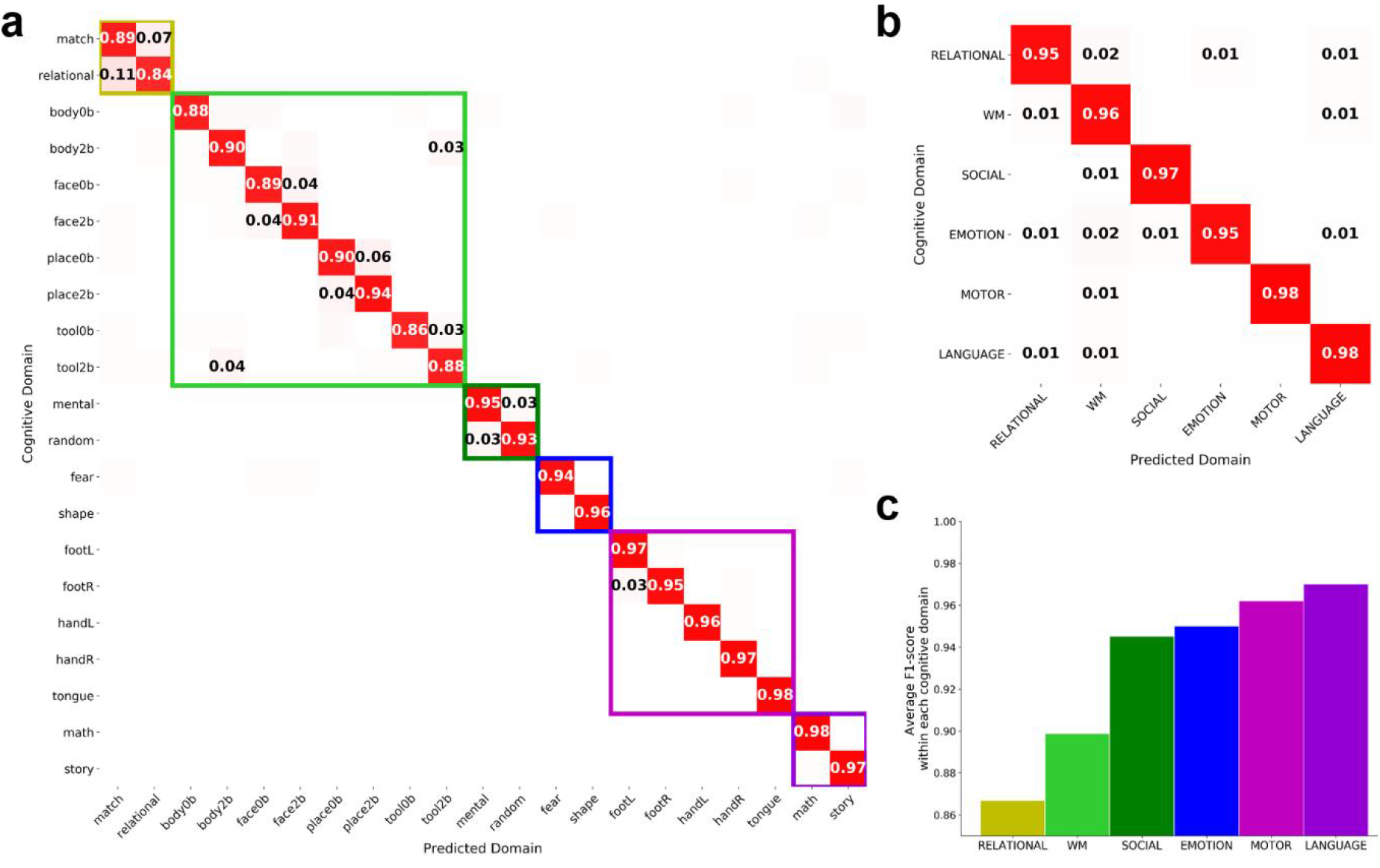
Confusion matrices of decoding 21 cognitive states and 6 cognitive domains. We calculated the confusion matrix of cognitive decoding for the predicting cognitive states using every 10s of fMRI signals. We used the ChebNet-K5 model on a functional graph derived from Glasser’s atlas in this analysis. The majority of misclassifications were found within the same cognitive domain rather than between domains. A clear block diagonal architecture of the confusion matrix, which indicates the majority of the cognitive tasks or domains were accurately identified, was shown for both 21 task conditions (**a**) and 6 cognitive domains (**b**). For the visualization purpose, a threshold of 0.02 was applied to the confusion matrix in (**a**), which indicates a low misclassification rate between domains (<2%). The averaged F1-score for each domain was shown in (**c**), which was calculated by averaging the scores within each cognitive domain based on the number of samples. Among the six cognitive domains, the Language tasks (in dark violet) and Motor tasks (in magenta) were the most easily recognizable conditions. Note: the six cognitive domains include relational processing (RELATIONAL), working memory (WM), social cognition (SOCIAL), facial emotional processing (EMOTION), body movements (MOTOR), and language comprehension (LANGUAGE).

### 3.3. Order *K* in ChebNet controls the propagation rate on the graph

At each ChebNet graph convolutional layer, the context of brain activity is propagated within a *K*-step neighborhood on the graph. As illustrated in Figure 2-S1, both the choices of K-order and graph architectures significantly impact the propagation rate of information flow in the brain, estimated by the expected first arrival time. We first investigated the impact of *K*-order on the decoding model by spanning over the list of [0,1,2,5,10], where K=0 indicates a global scaling factor on the input features by treating each node independently; K=1 indicates the integration of brain activity for each node and its direct neighbors, while K> 1 indicates the integration not limited to a region of interest or within a specific brain network, but instead expanding among inter-connected brain networks.

Our results (see Figure **3**) showed that the decoding performance gradually improved by increasing the *K*-order and reached a plateau when *K* ≥ 5. For example, when using a functional graph (Figure **3a**), the ChebNet- *K*1 model (i.e. ChebNet with *K* = 1) showed lower decoding accuracy than high-order models (91.67% vs 93.22%, respectively for *K* = 1 and *K* > 1), but was substantially higher than the *K* = 0 model (83.72%). When *K* ≥ 5, the training curves (indicated by the validation accuracy at the end of each training epoch) generally followed the same behaviors and the overall prediction accuracy plateaued approximately at the same level. We evaluated the sensitivity to the *K*- order for each cognitive domain, which demonstrated a strong domain-specific effect. For instance, when using functional-graph (Figure **3e**), the decoding on the Motor and Language tasks showed little impact by the *K*-order, which indicates that decoding on the two tasks was driven by functional interactions within the same functional network. On other hand, the decoding on the Working-memory and Relational-processing tasks showed high sensitivity to the choice of K-order and were continuously improved as increasing K, implying that the inter-network functional interactions might play an important role in the high-order cognitive functions. Similar findings were observed using the spatial-graph as well (Figure 3f), except that all cognitive tasks benefitted from high-order graph convolutions including Language tasks, which involve spatially distributed brain networks for language comprehension and numerical processing.

**Figure 3.**
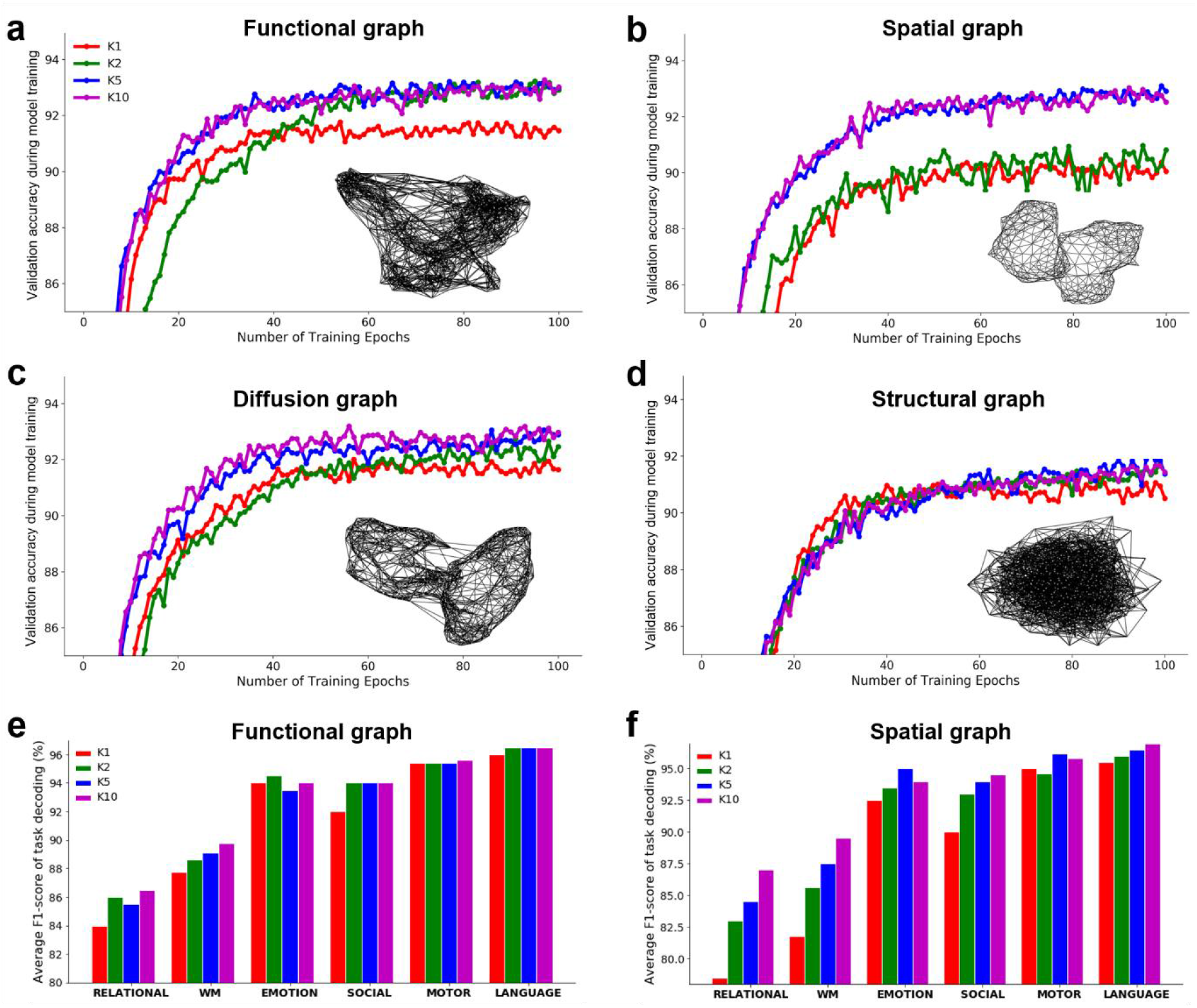
Effect of the *K*-order in ChebNet on four brain graphs and six cognitive domains. We evaluated four different types of brain graphs in the construction of ChebNet, including functional-graph computed using resting-state functional connectivity (**a**), spatial-graph representing spatial adjacency in the cortical surface (**b**), diffusion-graph estimated from whole-brain diffusion tractography (**c**), and structural-graph derived from structural covariance of cortical thickness across subjects (**d**). We first evaluated the effect of *K*-order on each brain graph in terms of the training curve, i.e. evaluating the model performance on the validation set at the end of each training epoch. We found a significant boost in decoding performance by using high-order graph convolutions (*K*>1) instead of integrating within the network (*K*=1) on all brain graphs. We also evaluated the impact of *K*-order on each cognitive domain by calculating the averaged F1-score on the test set and compared the effects across six cognitive domains and different brain graphs, e.g. the functional graph (**e**) and spatial graph (**f**). These results indicate that the decoding of high-order cognitive functions relies more on high-level interactions in the brain and this effect is more dramatic when using a graph architecture purely based on the topological organization of human brain.

### 3.4. Communities and brain networks accelerate information propagation on brain graph

There are multiple ways of capturing the network organization of human brain, using either spatial constraints (e.g. spatial adjacency in the geometry of brain surface (spatial-graph)), brain morphology (structural covariance of cortical thickness across a population of subjects (structural-graph)), or connectivity information (e.g. functional correlation of low-frequency fluctuations in BOLD signals (functional-graph) and tractography of fiber projections using diffusion-weighted images (diffusion-graph)), and others. Here, we mainly investigated these four types of brain graphs (Figure **4**). Among which, the spatial graph featured local geometric structures by connecting each parcel to its spatial neighbors on the cortical surface mesh. By contrast, modular and community structures have been widely demonstrated in the diffusion and functional brain graphs (Betzel and Bassett, 2017; Bullmore and Sporns, 2009). The structural graph (i.e. structural covariance of cortical thickness) somewhat resembles the functional graphs (Clos et al., 2014; Zielinski et al., 2010), but was contaminated by random noise effects (as shown in Figure **4c**).

**Figure 4.**
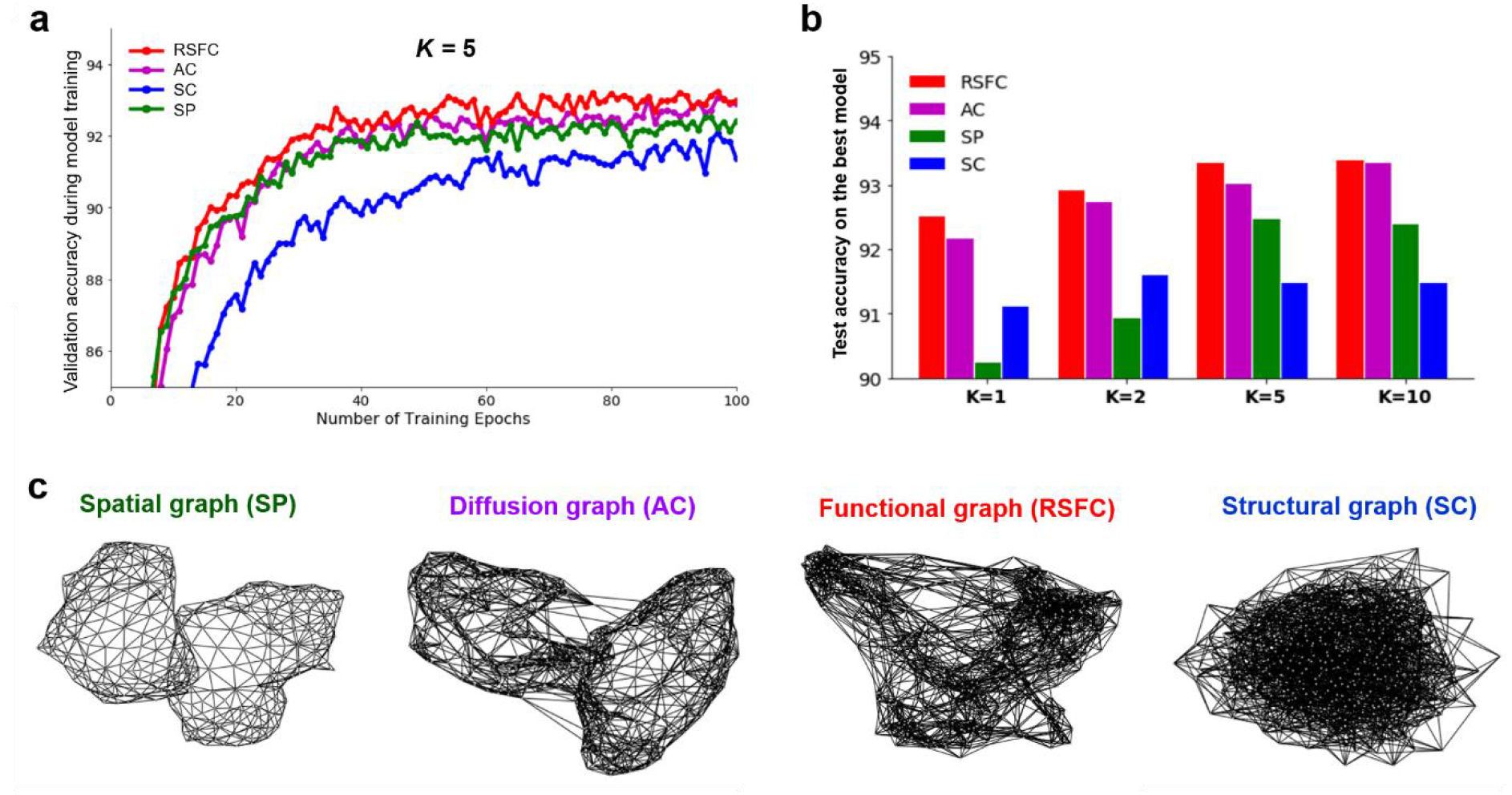
ChebNet decoding at different *K*-orders and using different brain graphs. We evaluated four different types of brain graphs for the multi-domain brain decoder, with the corresponding brain graph architecture shown in (**c**). We observed a strong interaction effect between brain graphs and *K*-orders (**b**) such that the decoding performance gradually improved by increasing the *K*-order for all brain graphs, but the graph architecture also had a big impact on the decoding performance. When fixing the *K*-order, for example using the ChebNet-*K*5 model (**a**), the functional (RSFC, in red) and diffusion graphs (AC, in purple) showed similar performance during model training, followed by the spatial graph (SP, in green) showing slightly lower decoding accuracy. The structural graph (SC, in blue) showed the lowest performance during the entire training process. Overall, the best decoding model was using ChebNet-*K*5 on the functional-graph (test accuracy=93.43%). To be noted that the baseline *K=0* model, which was invariant to the structure of brain graphs by treating each node independently, achieved much lower decoding performance on the same dataset (test accuracy=83.72%). Moreover, if we removed the graph convolutional layers from the decoding model, i.e. using a two-layer multilayer perceptron (MLP) with 256 and 64 neurons respectively, the decoding performance was further reduced (test accuracy=76.1%). These results indicate that high-order functional integration as well as the proper graph architectures are critical steps towards high-performing cognitive decoding.

The training curves on all brain graphs followed a similar trend as increasing the *K*-order (see Figure **3 a-d**) and showed similar behaviors when using a high propagation rate (i.e. K=5, as shown in Figure **4a**), except for the structural-graph which showed slightly lower performance than other graphs. Besides, we observed a significant interaction between the *K*-order and choices of brain graphs, such that the best decoding accuracy, evaluated on the test set after the entire training process, was achieved on the functional graph with a high-order model (Figure **4b**). When using spatial-graph (also shown in Figure **3b**), a gap in the decoding performance was detected between the low and high propagation groups (90.36% vs 92.4% for K≤2 and K>2, respectively) and reached a stable range after K>5 (93%). A much smaller impact of the K-order was shown in the functional-graph (Figure **3a**), with a smaller gap in brain decoding appearing at K=1 (92.52% vs 93.22%, respectively for K=1 and K>1). Diffusion-graph showed very similar behaviors as the functional-graph at all propagation rates (92.19% vs 93.04%, respectively for K=1 and K>1) and eventually converged to the same level of decoding performance when using a sufficiently high propagation rate (Figure **3c**). This is probably because the spatial-graph is only composed of short-distance connections, which restricts the integration of brain activity to a small local area. In order to reach out for the spatially distributed functional networks and even multiple brain systems, the model requires a relatively high K-order or a high path length, i.e. taking multiple walks on the graph. However, even with a high-order model, the spatial-graph still showed lower decoding performance compared to the diffusion and functional graphs (Figure **4b**). Our results indicated that the connectome-constrained graph architecture accelerates the propagation of information flow in the brain by integrating the neural dynamics through long-range connections. Moreover, considering that the sparsification of graph is the key to superior performance on graph learning benchmarks (Ye and Ji, 2021), we investigated functional graphs with different sparsity levels in brain decoding and found that, compared the original densely-connected brain connectomes, highly sparsified graphs performed much better at all K-orders (Fig.2- S2).

### 3.5. Functional homogeneity and small parcel size promotes local information processing

Another factor that significantly impacts the decoding model is the scale of the graph, i.e. number of brain parcels. Generally speaking, finer-scale atlases (smaller parcel size) will have higher internal homogeneity (Figure 5-S2 d) and result in less information loss after data projecting and averaging. Consequently, the decoding model using smaller parcels achieves better prediction of cognitive states and eventually reaches the plateau with a balance of model complexity and local homogeneity (Figure 5-S1 and S2). In order to investigate this effect, we have tested four different brain atlases derived from different modalities or human connectomes, including Yeo’s 7 and 17 functional networks (Yeo et al., 2011) (50 and 112 spatially continuous brain parcels), Brainnetome atlas (Fan et al., 2016) derived from diffusion tractography (246 brain parcels), and Glasser’s multi-model atlas (Glasser et al., 2016) (360 brain parcels), as well as brain atlases at multiple resolutions, e.g. Schaefer’s multiresolution brain parcellation (consisting of 100, 200, 400 and 1000 parcels respectively) (Schaefer et al., 2018). For each of the chosen atlases, we constructed the functional graph by calculating the resting-state functional connectivity based on 1080 subjects from the HCP database and evaluated the functional homogeneity of each brain parcel by calculating pairwise correlations of the connectivity patterns between all vertex within a parcel (Schaefer et al., 2018; Urchs et al., 2019). Coinciding with the literature that smaller parcels have higher functional homogeneity, we further demonstrated that finer-scale atlas results in higher decoding performance across different parcellation schemes (Figure 4-S2). Our results indicated that smaller parcel size resulted in higher functional homogeneity and better decoding performance (Figure 5-S2). As shown in Figure **5a**, the decoding performance started with a relatively low accuracy, for instance, 80.78% when using 50 spatially continuous regions derived from Yeo’s 7 network, and quickly improved by simply increasing the resolution of brain atlas, e.g. 85.23% when using 112 regions derived from the 17 network. Small improvement was detected when using around 300 regions or more (92.16% vs 93.43%, respectively for the Brainnetome atlas and Glasser’s atlas). Moreover, we observed that, on all brain atlases, the decoding performance gradually improved as increasing the *K*-order and reached the stable performance at *K*=5 (Figure **5b**).

**Figure 5.**
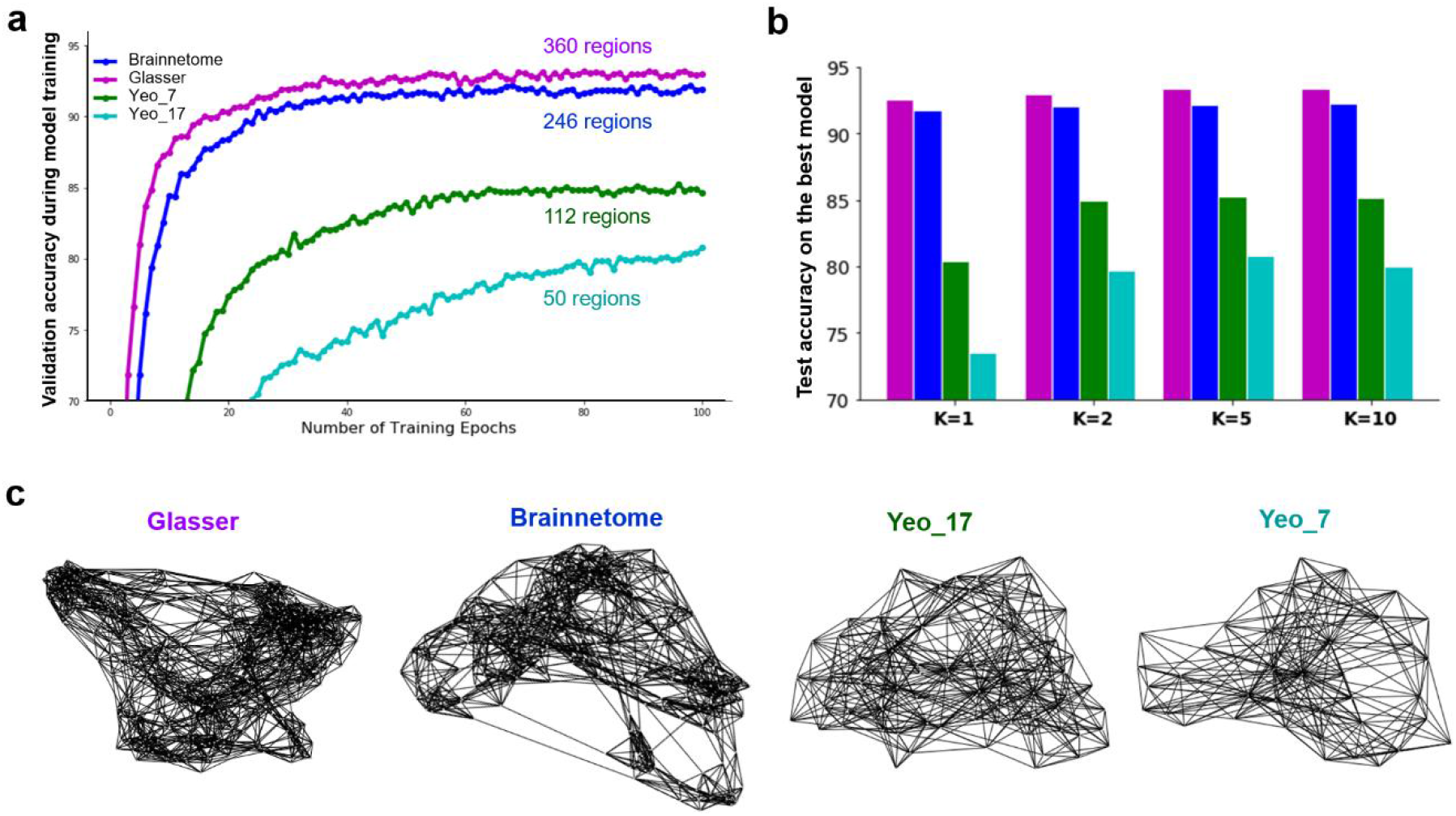
ChebNet decoding at *K*-orders and using different brain atlases. We evaluated different brain atlases at various resolutions for the construction of brain graph, including Yeo’s 7 and 17 functional networks (50 and 112 spatially continuous brain parcels, respectively), Brainnetome atlas derived from diffusion tractography (246 brain parcels), and Glasser’s multi-model atlas (360 brain parcels). For each brain atlas, we constructed the functional graph by evaluating the region-wise functional connectivity based on resting-state fMRI data (**c**). We trained separated decoding models at different *K*-orders and found that the Glasser’s atlas achieved the best decoding performance, followed by the Brainnetome atlas, and Yeo’s 17 and 7 functional atlases (**b**). When fixing the *K*-order, for example using the ChebNet-*K*5 model (**a**), we found significant advantages in model training by using finer-scale brain atlases (i.e. more brain regions and smaller parcel size) regardless of the information used for brain parcellation, e.g. functional, diffusion or multi-modal human connectomes.

We also evaluated Schaefer’s multiresolution brain parcellation (Schaefer et al., 2018), consisting of 100, 200, 400 and 1000 parcels respectively. As shown in Figure **6**, we found that higher resolution (smaller parcel size) improved the decoding performance at all *K*-orders and more dramatic improvement was detected at a small *K*-order than high-order models (e.g. K=1 vs K=5 in A and B, respectively). For instance, when using the ChebNet-K5 model, the decoding performance plateaued at 400 parcels with no further improvement by using higher resolutions, e.g. 1000 parcels. It’s worth noting that, compared to the Glasser’s atlas, the Schaefer’s atlas showed slightly lower performance at all resolutions and *K*-orders (e.g. 91.35% vs 93.43%, respectively for Schaefer’s (400 parcels) and Glasser’s atlas (360 parcels)). This is probably due to imperfect matching of cortical surface templates between different populations and consequently having much lower functional homogeneity when projecting Schaefer atlas onto the surface of HCP subjects (Figure 5-S1). The relationship of parcel size, functional homogeneity, and decoding accuracy was systematically investigated in Figure 5-S1. Our results indicate that the size and functional homogeneity of brain parcels significantly impact the decoding model. By contrast, how the brain atlas was constructed, for instance, using different imaging modalities or various types of connectome information, show little impact.

**Figure 6.**
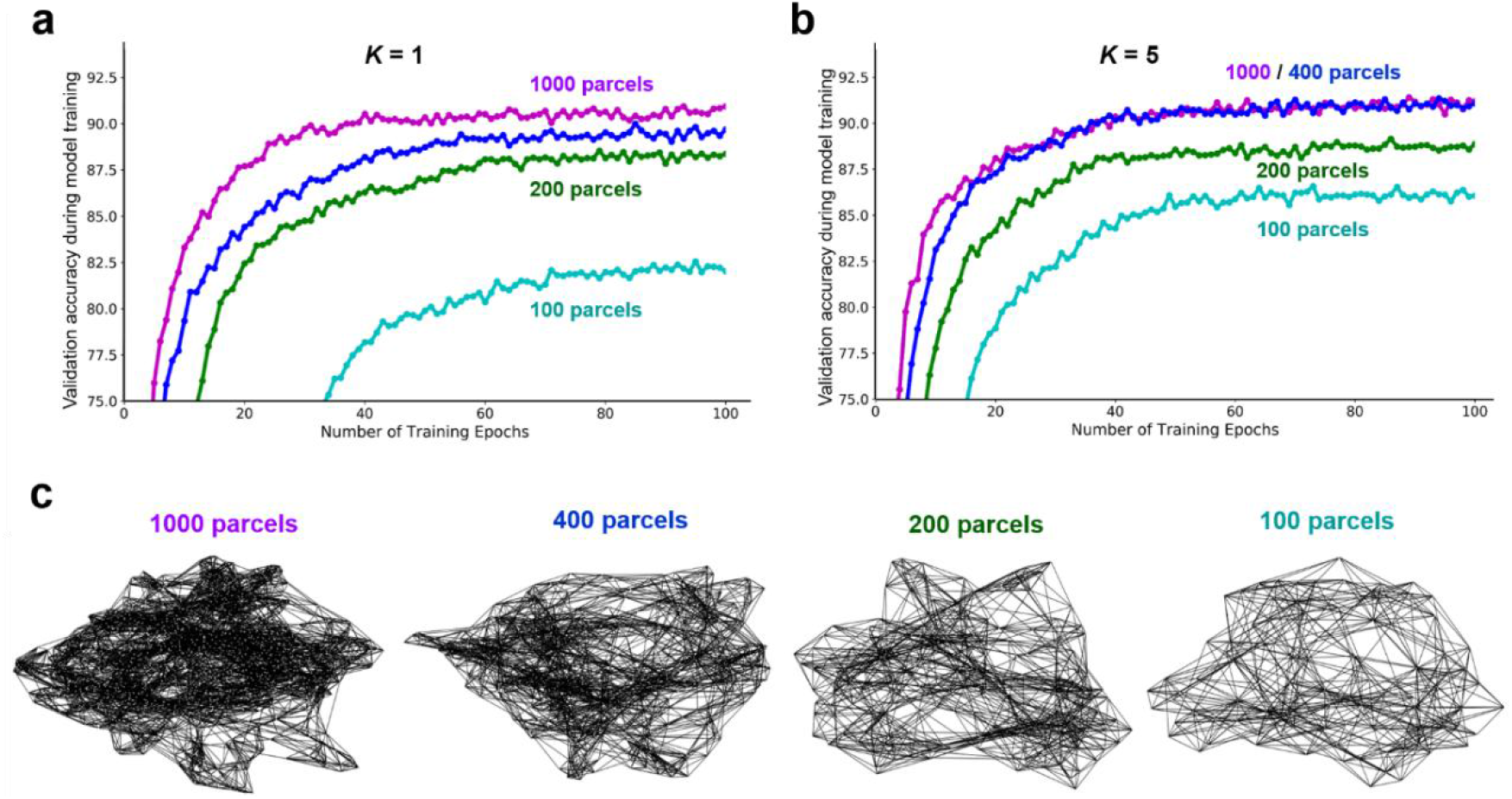
Brain decoding using functional brain parcellation at multiple resolutions. We evaluated different brain atlases derived from Schaefer’s multiresolution brain parcellation (Schaefer et al., 2018), consisting of 100, 200, 400 and 1000 brain parcels respectively. For each parcellation scheme, we constructed a functional graph by evaluating the region-wise functional connectivity based on resting-state fMRI data (c). We found that brain parcellation at higher resolutions (smaller parcel size or more regions) improved the performance of cognitive decoding, especially at a small K-order (e.g. K=1 in a). By contrast, when using a high-order model (e.g. K=5 in b), the model plateaued at 400 parcels with no further improvement on higher resolutions. However, we still observed improvements at small resolutions. It’s worth noting that, compared to the Glasser’s and Brainnetome atlases, both of which defined based on HCP subjects, the Schaefer’s atlas (derived from a different population) showed slightly lower performance (91.11% vs 92.16% vs 93.43%, respectively for Schaefer’s (400 parcels), Brainnetome (246 parcels) and Glasser’s atlas (360 parcels) when using the ChebNet-K5 model), probably due to imperfectly matching between surface templates across populations which consequently impacts the functional homogeneity of brain parcels (see Figure 5-S2).

### 3.6. High-order ChebNet adapts to network misspecification

The robustness of the decoding model was first evaluated by using randomized brain graphs (as shown in Figure **7c**), which were generated by randomly swapping a proportion of edges in the functional graph while keeping the node degrees unchanged. This procedure has been widely used as null models in network neuroscience literature, for instance (Betzel and Bassett, 2017; Sporns, 2018). Here we found that the network misspecifications only impact ChebNet decoding models with small Ks but barely influence the decoding performance when using high-order graph convolutions (Figure 7b). Specifically, when using a low random ratio (e.g. 0.1), the model achieved very similar decoding performance as the original functional-graph when *K* ≥ 2 (93.15% vs 93.22%, respectively for the randomized and functional graphs). When using a relatively high random ratio (e.g. 0.5), the decoding performance started with a low prediction accuracy and gradually improved by increasing the *K*-order (86.95% vs 91.52%, respectively for *K* < 2 and *K* ≥ 2), consistently lower than the original functional-graph at all *K*-orders. Besides, during model training, the randomized graphs showed much lower convergence speed than the original functional-graph and were prone to getting stuck in local minima, especially when only a small set of subjects were available (as shown in Figure 7-S1). Moreover, as stated before, structural-graph also exhibits some random effects (Figure **3d**), which resembles the behaviors of random graphs with the random ratio between 0.1 to 0.2 (91.13% vs 91.53%, respectively for *K* < 2 and *K* ≥ 2).

**Figure 7.**
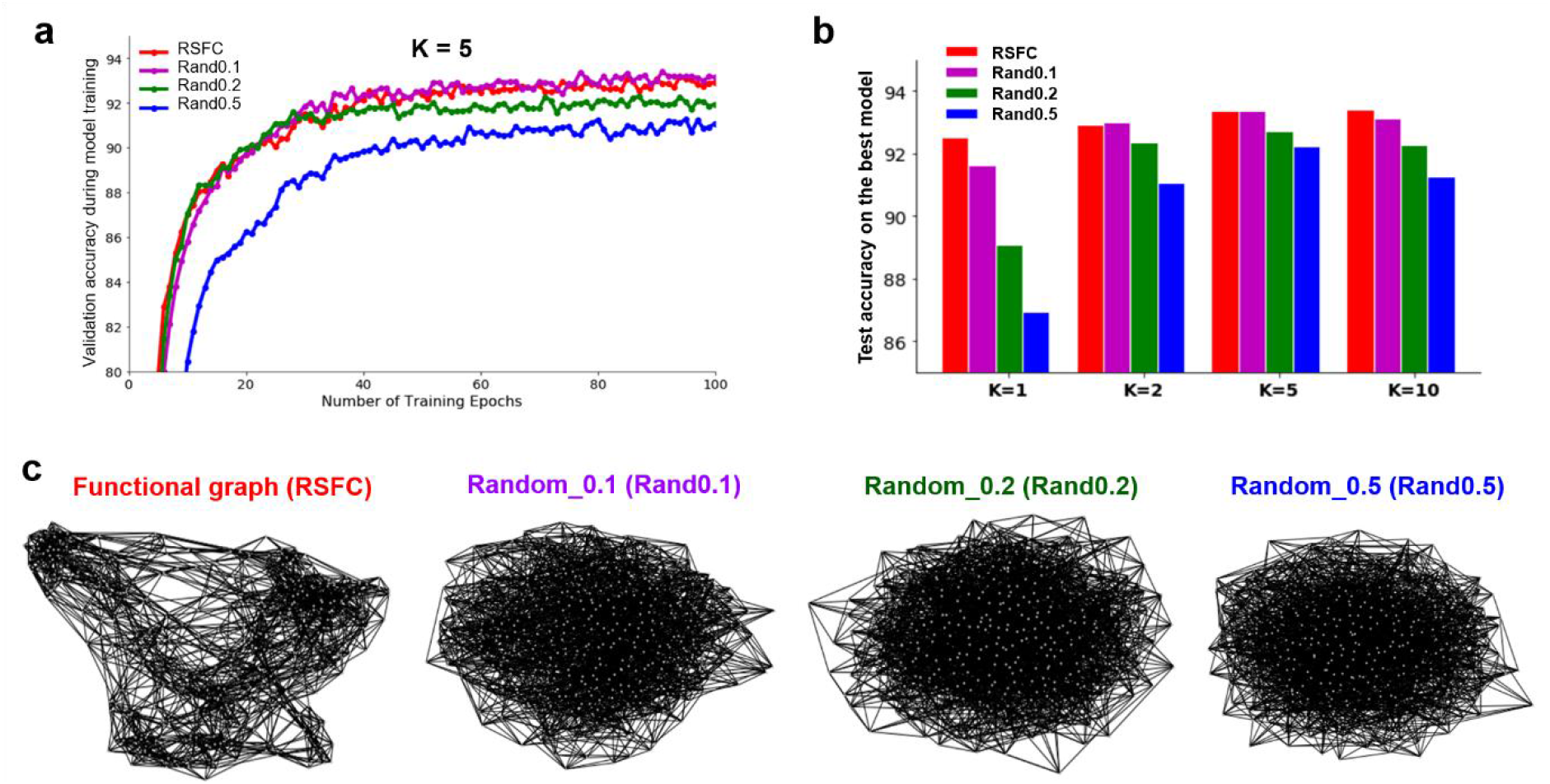
ChebNet decoding using random graphs derived from the functional graph. We generated three sets of randomized brain graphs by randomly swapping a portion of edges in the functional graph while keeping the node degree unchanged. The graph structures at different random ratios (e.g. 0.1, 0.2, 0.5) were shown in (**c**). When fixing the *K*-order, for example using the ChebNet- *K*5 model (**a**), the functional graph (RSFC, in red) and its low randomness copy (Rand0.1, in purple) showed very similar performance during model training, followed by the moderately (Rand0.2, in green) and highly randomized graph (Rand0.5, in blue). We observed a strong interaction effect between randomness ratios and the *K*-orders (**b**) such that the decoding performance on randomized graphs started with a low prediction accuracy and gradually improved when increasing the *K*-orders, consistently lower than the original functional-graph at all *K*-orders. The impact of graph randomization was more dramatic at a small *K*-order (e.g. *K*=1) and smaller impact was detected when using high-order models. The best performance was achieved by using a high-order model on the functional graph or its low random copies.

### 3.7. Robustness to random attacks in brain regions and networks

The robustness of the decoding model was also evaluated under random attacks, by removing brain responses from a small set of regions in the brain. This procedure has been commonly used to simulate brain lesions due to brain injury or neurological disorders (Alstott et al., 2009; Honey and Sporns, 2008). Here, we mainly focused on two types of attacks, either using randomly chosen regions or targeted regions from a specific brain network. Our results demonstrated that the proposed ChebNet graph convolution was resilient to random attacks on regions but not networks. Specifically, lesions on a small set of randomly chosen brain regions, e.g. 40 parcels or less, did not affect the decoding performance (median decay in the decoding accuracy: 1.2%). More severe decays were detected as more regions were affected by lesions (median decay: 2.8%, 15.3% and 40.4% respectively for 72, 180 and 288 affected regions). This pattern of performance decay was not related to the betweenness or centrality of targeted regions (r=0.091, p=0.08), but rather associated with their involvement in task-related brain activations. For instance, the decay in the decoding of WM tasks was mostly driven by areas in the ventral visual stream, including PH, VVC and V8, as well as frontoparietal network regions, including PFm, MFG and area 6r, as revealed by a correlation analysis between the decay of decoding performance and the infection of each brain region (as shown in Figure **8a**). These regions have been reported to be engaged in the process of WM tasks (Christophel et al., 2012; Harrison and Tong, 2009; Mayer et al., 2007) with a similar pattern revealed by the saliency map analysis (as shown in Figure **9**). On the other hand, much larger decays were observed when switching to network attacks (around 10%), with most severe cases found in the limbic, frontoparietal network (FPN) and attention networks (consisting of 60, 82, 104 affected parcels, respectively). A similar pattern of decays was observed when using the 17-network parcellation, with the largest decays found in the networks related to cognitive control and attention (as shown in Figure **8b** and **8c**). Interestingly, the distribution of performance decays in cognitive decoding was not related to the size of affected intrinsic networks (r=0.097, p=0.332). To conclude, the ChebNet decoding model was robust to random attacks but was more vulnerable to targeted attacks on task-related brain regions and networks. Moreover, compared to the low-order model (e.g. K=1), the high-order ChebNet model was more resilient to random or targeted attacks by showing smaller decays in cognitive decoding due to lesions on regions and networks (Figure 8-S1).

**Figure 8.**
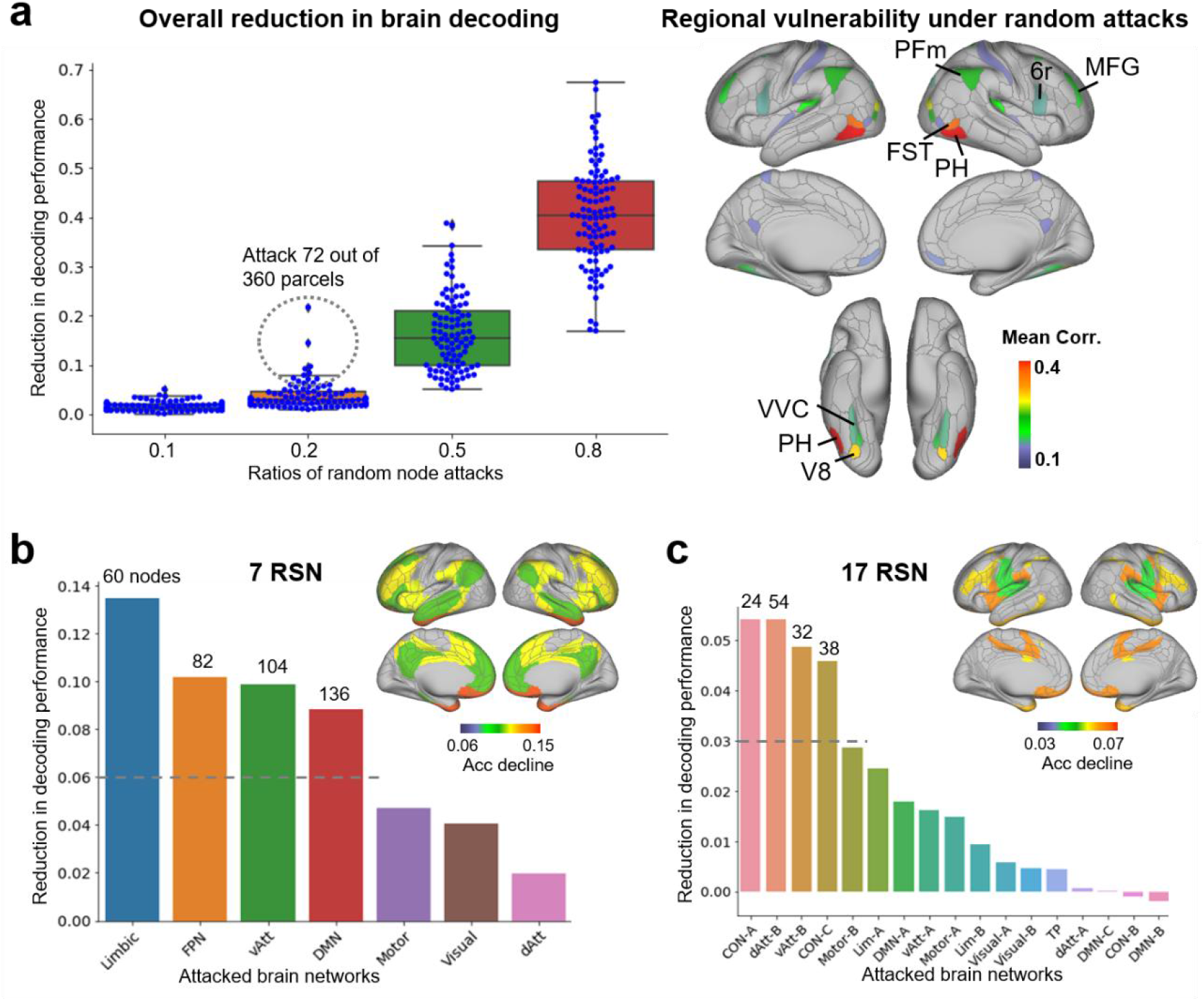
Robustness of ChebNet decoding models to random attacks on nodes and networks. We simulated two types of brain lesions, either attacking randomly chosen regions or targeting for regions from a specific brain network. For random node attacks (**a**), we silenced brain responses from randomly chosen nodes, ranging from 10% to 80% of brain parcels, and repeated the process for 100 times. We found that the decoding model was highly resilient to random node attacks by showing a small decay in the decoding accuracy (<3% when attacking around 80 regions). The most vulnerable regions include areas in the ventral and dorsal visual streams (top right panel). For network attacks, we removed one brain network at a time, identified by Yeo’s 7 (**b**) and 17 (**c**) resting-state functional networks, and silenced the associated brain parcels with at least 30% areas affected. We found severe decays in the cognitive decoding when attacking specific brain networks (around 10% and 5% for 7- and 17-network parcellation, respectively). Among which, the most vulnerable networks were those related to cognitive control and attention, for instance the frontoparietal network (FPN) and ventral attention networks (vAtt).

**Figure 9.**
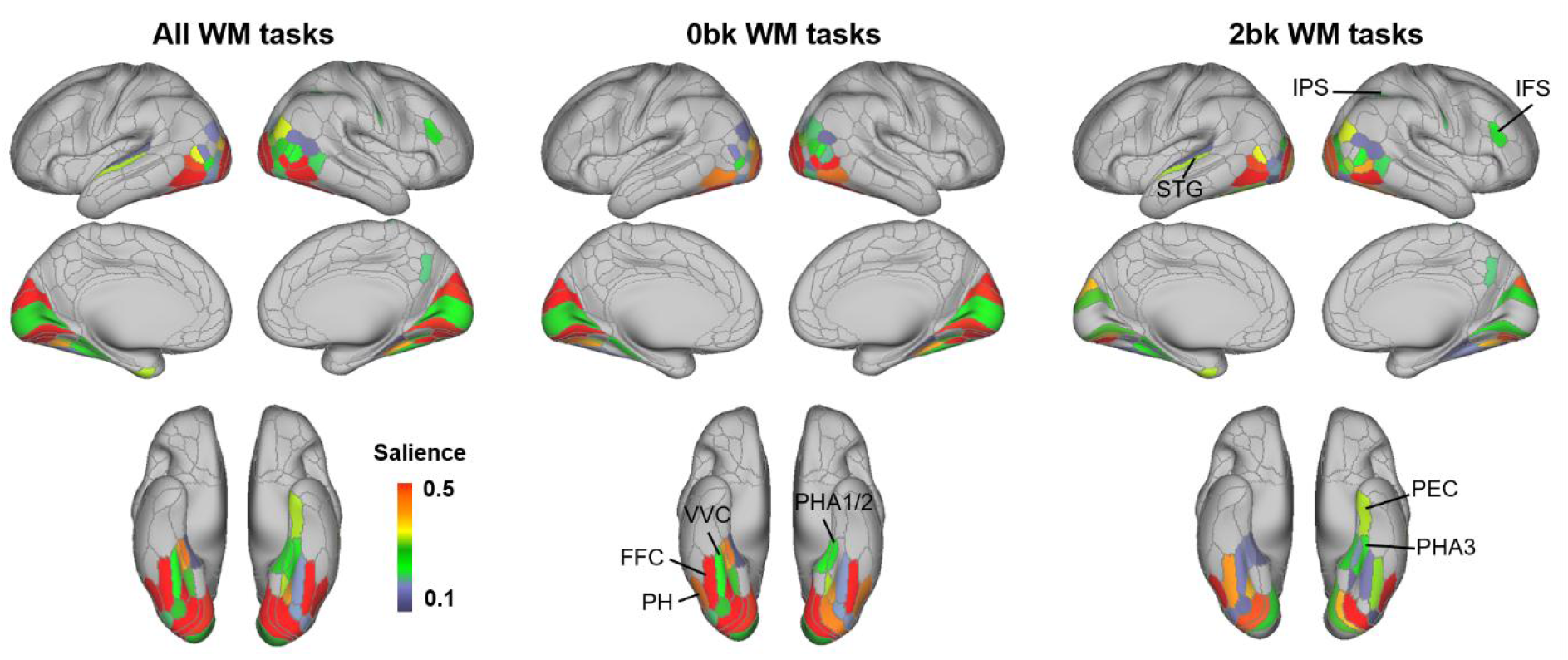
Saliency maps for the decoding of WM tasks. We generated the salient features that showed high contributions to the classification of WM tasks by using the guided backpropagation approach. The model detected biologically meaningful and category-specific salient features on 0back and 2back WM tasks. For 0back tasks, the model identified salient regions in the visual cortex and ventral visual stream, including V1-V3, PH, FFC and PHA. For 2back tasks, the model detected salient features in the frontoparietal network regions, including IPS and MFG, and the temporal regions, e.g. superior temporal gyrus (STG) and Perirhinal Cortex (PRC). Note that only brain regions with a high saliency (saliency values>0.1, full range of saliency is (0,1)) and a significant ‘task condition’ effect (p<0.001) were shown in the brain maps.

### 3.8. Node contributions revealed by saliency maps

In order to illustrate the biological basis of the decoding model, we generated the saliency maps on the trained model by propagating the non-negative gradients backwards to the input layer (Springenberg et al., 2014). An input feature is *salient* or *important* only if its little variation causes big changes in the decoding output. As shown in Figure **9**, category-specific salient features were detected for all WM tasks and separately for 0back and 2back conditions. For 0back tasks, regions in the visual cortex and ventral visual stream were identified, including V1-V3, PH, FFC and PHA, coinciding with the literature that the ventral visual stream is involved in the recognition of visual stimuli (Haan and Cowey, 2011; Haxby et al., 2014). By contrast, the frontoparietal network regions, including IPS and MFG, and the temporal regions, e.g. superior temporal gyrus (STG) and Perirhinal Cortex (PRC), were contributed to the decoding of 2back tasks. These regions have been shown to be engaged in object recognition and visual working memory (Eriksson et al., 2015; Schon et al., 2016). Our findings revealed that the ChebNet decoding model captured task-related brain regions and category-specific patterns of brain activity, and consequently extracted biologically meaningful features for the classification of different task conditions. This analysis also revealed high salience in the primary sensorimotor cortices for Motor tasks (Fig.9-S1a) and in the perisylvian language-related brain regions for Language tasks (Fig.9-S1b). Moreover, the learned graph representations, i.e. activation maps in the last ChebNet layer, demonstrated distinct patterns of brain activity in the perisylvian language-related regions among mathematics and story trials (Fig.9-S1c).

## 4. Discussion

We proposed a generalized framework for brain decoding based on ChebNet graph convolutions. The model takes in a short window of fMRI time series and a brain graph (with nodes representing brain parcels and edges representing brain connectivity), and integrates brain activity in a multiscale manner, ranging from local regions, to brain circuits/networks, and across multiple brain systems. Using a 10s- time window of fMRI signals, our model identified 21 different task conditions across multiple cognitive domains with a test accuracy of 93% (chance level of 4.8%). We systematically investigated the impacts of using high-order graph convolutions and connectome-constrained graph architectures in the decoding model. Our findings revealed that 1) connectome constraints accelerate the propagation and integration of brain activity by adding shortcuts through long-range connections; 2) smaller parcel size or finer-scale atlas improves the internal homogeneity of brain parcels and boosts the overall decoding performance; 3) high-order graph convolution encodes functional integration of distributed brain activity at multiple scales and boots the decoding performance on all brain graphs. These results support an important role of functional integration in cognitive decoding, which leads to higher decoding accuracy as well as better adaptation to network misspecifications due to random rewiring and node attacks.

### 4.1. Functional integration plays an important role in cognitive decoding

A variety of computational models have been proposed in the field of brain decoding in the last decades, with the aim of learning a linear discriminative function on the spatial patterns of brain activation under specific experimental conditions. For instance, researchers have successfully attempted to use brain activity to reconstruct the frames of movies (Nishimoto et al., 2011), or to decode the semantic context from words (Mitchell et al., 2008) and visual scenes (Huth et al., 2012) by using linear regression models. Recently, with the fast development of deep learning, artificial neural networks have been applied in the field of decoding human cognition from recorded brain activity, for instance using classical convolutional (Wang et al., 2020) and recurrent neural networks (Li and Fan, 2019), or a generalized form of convolutions in the graph domain (Zhang et al., 2021). However, the majority of current brain decoding studies so far focused on segregating neural substrates of different cognitive tasks by treating each brain area independently in the decoding model (Haxby, 2012; Haxby et al., 2014; Poldrack, 2011; Varoquaux et al., 2018). The interactions between different brain areas and networks have been largely ignored, but starts to play an important part in the field. Cole and colleagues (Cole et al., 2016) demonstrated the possibility of predicting activations of unseen brain regions or new subjects by transmitting information flow of brain activity within a functional network. Using a similar idea, Li and Fan (Li and Fan, 2019) have successfully inferred cognitive states from network-level neural activity by integrating signals from large-scale brain networks. Accordingly, we recently proposed a GNN architecture that propagates brain activity through functional connectomes and integrates the context of brain activity in both local and global extent (Zhang et al., 2021). The model has achieved high decoding accuracy over a variety of cognitive domains −90% in (Zhang et al., 2021)- outperforming other baseline approaches on the benchmark, including non-integrative full-brain models, e.g. 64% in linear SVM and 76% in MLP.

In this study, we further extend the GNN framework by enlarging the scale of functional integration in graph convolutions and exploring the impacts of different connectome priors in cognitive decoding. Firstly, we implement high-level integration of task-evoked brain activity in the decoding model, extending from a single brain network (K=1, similar to (Zhang et al., 2021)), to multiple brain systems (K>1) and towards the full brain (K>5). Secondly, we explore different choices of graph architectures that restrict the propagation of brain activity through the topology or connectome of the human brain. Our results demonstrate a significant improvement in large-scale cognitive decoding by implementing high-order interactions on functional connectomes (93% using the 5-order ChebNet model). We observed this effect on all types of brain graphs (Figure **3**). In addition, the high-order integrative model resulted in a similar level of decoding performance when using different connectome priors, for instance the diffusion-graph derived from whole-brain tractography of fiber projections (Figure **4**), but higher than topological and morphological priors.

The sensitivity to functional integration was not uniformly distributed among all cognitive domains but rather depended on the cognitive demands of tasks (Figure **3e**). For instance, WM and Relational-processing required a high-order model to integrate brain activity across multiple brain systems, while Language and Motor tasks already achieved the optimal performance by taking into account the interactions within targeted brain networks. These findings may partially explain the excellent decoding performance in previous decoding studies when tackling unimodal cognitive functions, for instance, distinguishing body movements using patterns of brain activity from in the motor cortex (Mottolese et al., 2013) or restricting the context of brain activity within a local area (Wang et al., 2020), but showed poor decoding on high-order cognitive functions, for instance, classifying 0-back and 2-back WM tasks (Wang et al., 2020). By contrast, when taking into account the integration within the functional networks, the model achieved high accuracy on both unimodal and high-order cognitive functions (Li and Fan, 2019; Zhang et al., 2021), and showed a further improvement by implementing high-order interactions among brain networks (Figure 3e). Our results indicate that high-performing cognitive decoding relies on the encoding of high-level functional integration in brain activity, not limited to local brain activity or low-level interactions. In addition to higher decoding accuracy, the integrative model also improved the robustness to perturbations on the graph architecture, for instance, by using different types of interactions (Figure **4**), network rewiring (Figure **7**), as well as random attacks on brain regions and networks (Figure **8**). These findings greatly expand current main perspectives on brain decoding that aims to segregate neural substrates of different cognitive tasks, and suggest an important role of functional integration during cognitive processes, especially for high-order cognitive functions.

### 4.2. Connectome-constrained graph architectures accelerate information propagation and functional integration

A series of studies in the literature have illustrated a great potential of using resting-state functional connectivity in predicting cognitive functions, not only in behavior (Rosenberg et al., 2020; Yamashita et al., 2018) but also in task-evoked brain activity (Cole et al., 2016; Tavor et al., 2016). Recent decoding studies have shown that the connectivity patterns (Richiardi et al., 2011; Shirer et al., 2012) as well as the activity flow on functional networks (Li and Fan, 2019; Zhang et al., 2021) are also predictive of cognitive states. In the current study, we further demonstrated the potentials of using other types of human connectomes beyond functional networks. As one of the basic graph architectures, brain topology and its derived spatial constraints has been widely used in fMRI analysis, for instance generating brain parcels on individual brains (Blumensath et al., 2013; Craddock et al., 2012; Ma et al., 2021). When implementing GNN with such constraints, the model generalizes the classical convolutional operations defined in the 3D volume space, e.g., (Wang et al., 2020), onto the geometry of cortical surfaces. Interestingly, the topologically constrained decoding model achieved a decent performance on unimodal cognitive functions, similar to previous studies (Wang et al., 2020), but showed lower performance on high-order cognitions compared to functional connectomes (Figure **3f**). The morphological brain networks, e.g. structural covariance of cortical thickness, on the other hand, resembled randomized functional connectomes by showing slightly lower prediction accuracy than the original graph in all cases (Figures **4** and **6**). Compared to them, the biologically constrained graph convolutions using either functional or diffusion brain connectomes achieved the highest decoding performance along with high robustness to network misspecifications due to random rewiring and node attacks. Our results revealed a clear gradient in the contribution of graph architectures for cognitive decoding, ranging from the lowest decoding performance by restricting activity flow within local areas (spatial-graph), to moderate performance by integrating information within neural circuits (diffusion-graph), and to the highest performance when using functional networks (functional-graph). Moreover, this gradient was only dominant at a low propagation rate and mostly diminished when using a high propagation rate, by converging to a similar level on all brain graphs (Figure **4**). We observed a similar interaction effect on the randomized functional graphs as well (Figure **7**). One possible explanation is that the spatial graph only consists of short-distance connections, which restricts information integration to a small local area on the cortical surface. When using a high-order propagation rate (K>1), the model expands the receptive field or neighborhood size at each graph convolutional layer by taking multiple steps at once, and eventually reaches a large neighborhood that includes distributed brain regions from a task-related functional network when *K* is sufficiently high (K>5). On the other hand, by adding modular and community structures to the graph, extracted from diffusion and functional connectomes, the model further accelerates the propagation of brain activity in the brain by adding shortcuts through long-range connections. Numerous studies have illustrated that the diffusion and functional brain networks, rather than regular spatial networks, followed the small-world configuration with information segregation and integration at low wiring and energy costs (Bullmore and Sporns, 2009; Bullmore and Bassett, 2011; Liao et al., 2017).

Our results revealed that, in addition to the K-order in ChebNet graph convolution, which controls the propagation rate of information flow on the graph by taking multiple walks at once, the architecture of brain graphs restricts the scale of the information flow at each step, ranging from local areas (spatial-graph), to neural circuits (diffusion-graph), and to functional networks (functional-graph). Among which, the biologically constrained graph structures derived from human brain connectomes highly boost the decoding of cognitive processes.

### 4.3. Limitations and Future directions

In this study, we only investigated the choices of group-wise brain graphs in the decoding model, i.e. all subjects taking the same definition of nodes and edges in the graph architecture. The benefit of using such a brain graph is that the decoding model can easily generalize across large populations (more than 1000 subjects in our case), transfer onto new populations and datasets, as well as better adaptation to network misspecifications due to random rewiring and node attacks. However, our findings did not rule out the possibility of using an optimal individual brain graph in cognitive decoding, for instance constructing an individualized brain parcellation and subject-level graph architecture. This might be beneficial when handling massive fMRI data from a single subject. Due to the limited amount of functional imaging data from individual brains in the HCP database, this option was not explored in the current study. In the next project, we plan to investigate how individual brain parcellation schemes and individual graph architectures improve brain decoding of complex cognitive processes on individual brains by using fMRI data from a local data collection (www.cneuromod.ca).

## 5. Conclusions

In summary, we propose a connectome-based graph neural network to encode the underlying spatiotemporal organization of cognitive processes and explore the optimal connectome architectures for large-scale cognitive decoding, including the path length, the homogeneity of brain parcels and the type of interactions. The model propagates information flow of brain dynamics in a multiscale manner, ranging from localized brain areas, to a specific brain network and towards the full brain. The scale of functional integration is largely controlled by the path length, i.e. K-order of ChebNet, and is restricted by the nature of interactions, i.e. brain connectomes. Compared to the topological and morphological constraints, connectome-constrained graph convolutions achieved better decoding of cognitive states. The edge-sparsified graph structures also contribute to superior performance of the ChebNet decoding model. Together, our findings indicate that human connectome constraints and multiscale functional integration are critical for large-scale cognitive decoding. Such decoding models better adapt to small fluctuations on the graph architecture including disconnections and lesions in the brain.

## Acknowledgment

This work was partially supported by the Science and Technology Innovation 2030 - Brain Science and Brain-Inspired Intelligence Project (Grant No. 2021ZD0200201), Scientific Project of Zhejiang Lab (No.2021ND0PI01, No.2022ND0AN01), Courtois foundation through the Courtois NeuroMod Project and the IVADO Postdoctoral Scholarships Program. PB is supported by a salary award of “Fonds de recherche du Québec - Santé”, chercheur boursier junior 2.

## Author contributions

Conceptualization: YZ, PB; Methodology: YZ, PB; Visualization: YZ, PB;

Investigation: YZ, NF, PB;

Writing—original draft: YZ, NF, PB

Writing—review & editing: YZ, NF, PB

## Data and materials availability

We use the block-design task-fMRI dataset from the Human Connectome Project S1200 release, downloaded from https://db.humanconnectome.org/data/projects/HCP_1200. In total, fMRI data from 1095 unique subjects under six different task domains and resting-state are used in this study. The minimal preprocessed fMRI data are used, in both CIFTI and NIFTI formats, depending on the choice of brain parcellation schemes. The proposed ChebNet decoding pipeline, as well as the optimized decoding models and the construction of brain graphs, are made publicly available in the following repository: https://github.com/zhangyu2ustc/gcn_tutorial_test.git.

## References

Alstott, J., Breakspear, M., Hagmann, P., Cammoun, L., Sporns, O., 2009. Modeling the Impact of Lesions in the Human Brain. PLOS Comput. Biol. 5, e1000408. https://doi.org/10.1371/journal.pcbi.1000408

Barch, D.M., Burgess, G.C., Harms, M.P., Petersen, S.E., Schlaggar, B.L., Corbetta, M., Glasser, M.F., Curtiss, S., Dixit, S., Feldt, C., Nolan, D., Bryant, E., Hartley, T., Footer, O., Bjork, J.M., Poldrack, R., Smith, S., Johansen-Berg, H., Snyder, A.Z., Van Essen, D.C., 2013. Function in the human connectome: Task-fMRI and individual differences in behavior. NeuroImage, Mapping the Connectome 80, 169–189. https://doi.org/10.1016/j.neuroimage.2013.05.033

Bassett, D.S., Bullmore, E.T., 2009. Human brain networks in health and disease. Curr. Opin. Neurol. 22, 340–347. https://doi.org/10.1097/WCO.0b013e32832d93dd

Betzel, R.F., Bassett, D.S., 2017. Multi-scale brain networks. NeuroImage, Functional Architecture of the Brain 160, 73–83. https://doi.org/10.1016/j.neuroimage.2016.11.006

Blumensath, T., Jbabdi, S., Glasser, M.F., Van Essen, D.C., Ugurbil, K., Behrens, T.E.J., Smith, S.M., 2013. Spatially constrained hierarchical parcellation of the brain with resting-state fMRI. NeuroImage 76, 313–324. https://doi.org/10.1016/j.neuroimage.2013.03.024

Bullmore, E., Sporns, O., 2009. Complex brain networks: graph theoretical analysis of structural and functional systems. Nat. Rev. Neurosci. 10, 186–198. https://doi.org/10.1038/nrn2575

Bullmore, E.T., Bassett, D.S., 2011. Brain graphs: graphical models of the human brain connectome. Annu. Rev. Clin. Psychol. 7, 113–140. https://doi.org/10.1146/annurev-clinpsy-040510-143934

Christophel, T.B., Hebart, M.N., Haynes, J.-D., 2012. Decoding the Contents of Visual Short-Term Memory from Human Visual and Parietal Cortex. J. Neurosci. 32, 12983–12989. https://doi.org/10.1523/JNEUROSCI.0184-12.2012

Christophel, T.B., Klink, P.C., Spitzer, B., Roelfsema, P.R., Haynes, J.-D., 2017. The Distributed Nature of Working Memory. Trends Cogn. Sci. 21, 111–124. https://doi.org/10.1016/j.tics.2016.12.007

Clos, M., Rottschy, C., Laird, A.R., Fox, P.T., Eickhoff, S.B., 2014. Comparison of structural covariance with functional connectivity approaches exemplified by an investigation of the left anterior insula. NeuroImage 99, 269–280. https://doi.org/10.1016/j.neuroimage.2014.05.030

Cole, M.W., Ito, T., Bassett, D.S., Schultz, D.H., 2016. Activity flow over resting-state networks shapes cognitive task activations. Nat. Neurosci. 12.

Craddock, R.C., James, G.A., Holtzheimer, P.E., Hu, X.P., Mayberg, H.S., 2012. A whole brain fMRI atlas generated via spatially constrained spectral clustering. Hum. Brain Mapp. 33, 1914–1928. https://doi.org/10.1002/hbm.21333

Eickhoff, S.B., Yeo, B.T.T., Genon, S., 2018. Imaging-based parcellations of the human brain. Nat. Rev. Neurosci. 19, 672–686. https://doi.org/10.1038/s41583-018-0071-7

Eriksson, J., Vogel, E.K., Lansner, A., Bergström, F., Nyberg, L., 2015. Neurocognitive Architecture of Working Memory. Neuron 88, 33–46. https://doi.org/10.1016/j.neuron.2015.09.020

Fan, L., Li, H., Zhuo, J., Zhang, Y., Wang, J., Chen, L., Yang, Z., Chu, C., Xie, S., Laird, A.R., Fox, P.T., Eickhoff, S.B., Yu, C., Jiang, T., 2016. The Human Brainnetome Atlas: A New Brain Atlas Based on Connectional Architecture. Cereb. Cortex 26, 3508–3526. https://doi.org/10.1093/cercor/bhw157

Glasser, M.F., Coalson, T.S., Robinson, E.C., Hacker, C.D., Harwell, J., Yacoub, E., Ugurbil, K., Andersson, J., Beckmann, C.F., Jenkinson, M., Smith, S.M., Van Essen, D.C., 2016. A multi-modal parcellation of human cerebral cortex. Nature 536, 171–178. https://doi.org/10.1038/nature18933

Glasser, M.F., Sotiropoulos, S.N., Wilson, J.A., Coalson, T.S., Fischl, B., Andersson, J.L., Xu, J., Jbabdi, S., Webster, M., Polimeni, J.R., Van Essen, D.C., Jenkinson, M., 2013. The minimal preprocessing pipelines for the Human Connectome Project. NeuroImage, Mapping the Connectome 80, 105–124. https://doi.org/10.1016/j.neuroimage.2013.04.127

Haan, E.H.F. de, Cowey, A., 2011. On the usefulness of ‘what’ and ‘where’ pathways in vision. Trends Cogn. Sci. 15, 460–466. https://doi.org/10.1016/j.tics.2011.08.005

Harrison, S.A., Tong, F., 2009. Decoding reveals the contents of visual working memory in early visual areas. Nature 458, 632–635. https://doi.org/10.1038/nature07832

Haxby, J.V., 2012. Multivariate pattern analysis of fMRI: The early beginnings. NeuroImage 62, 852–855. https://doi.org/10.1016/j.neuroimage.2012.03.016

Haxby, J.V., Connolly, A.C., Guntupalli, J.S., 2014. Decoding Neural Representational Spaces Using Multivariate Pattern Analysis. Annu. Rev. Neurosci. 37, 435–456. https://doi.org/10.1146/annurev-neuro-062012-170325

Haxby, J.V., Gobbini, M.I., Furey, M.L., Ishai, A., Schouten, J.L., Pietrini, P., 2001. Distributed and overlapping representations of faces and objects in ventral temporal cortex. Science 293, 2425–2430. https://doi.org/10.1126/science.1063736

Hearne, L.J., Mill, R.D., Keane, B.P., Repovš, G., Anticevic, A., Cole, M.W., 2021. Activity flow underlying abnormalities in brain activations and cognition in schizophrenia. Sci. Adv. 7, eabf2513. https://doi.org/10.1126/sciadv.abf2513

Honey, C.J., Sporns, O., 2008. Dynamical consequences of lesions in cortical networks. Hum. Brain Mapp. 29, 802–809. https://doi.org/10.1002/hbm.20579

Huth, A.G., Nishimoto, S., Vu, A.T., Gallant, J.L., 2012. A continuous semantic space describes the representation of thousands of object and action categories across the human brain. Neuron 76, 1210–1224. https://doi.org/10.1016/j.neuron.2012.10.014

Ito, T., Hearne, L., Mill, R., Cocuzza, C., Cole, M.W., 2020. Discovering the Computational Relevance of Brain Network Organization. Trends Cogn. Sci. 24, 25–38. https://doi.org/10.1016/j.tics.2019.10.005

Li, H., Fan, Y., 2019. Interpretable, highly accurate brain decoding of subtly distinct brain states from functional MRI using intrinsic functional networks and long short-term memory recurrent neural networks. NeuroImage 202, 116059. https://doi.org/10.1016/j.neuroimage.2019.116059

Liao, X., Vasilakos, A.V., He, Y., 2017. Small-world human brain networks: Perspectives and challenges. Neurosci. Biobehav. Rev. 77, 286–300. https://doi.org/10.1016/j.neubiorev.2017.03.018

Ma, L., Zhang, Y., Zhang, H., Cheng, L., Zhuo, J., Shi, W., Lu, Y., Li, W., Yang, Z., Wang, J., Fan, L., Jiang, T., 2021. BAI-Net: Individualized Human Cerebral Cartography using Graph Convolutional Network. https://doi.org/10.1101/2021.07.15.452577

Margulies, D.S., Ghosh, S.S., Goulas, A., Falkiewicz, M., Huntenburg, J.M., Langs, G., Bezgin, G., Eickhoff, S.B., Castellanos, F.X., Petrides, M., Jefferies, E., Smallwood, J., 2016. Situating the default-mode network along a principal gradient of macroscale cortical organization. Proc. Natl. Acad. Sci. 113, 12574–12579. https://doi.org/10.1073/pnas.1608282113

Mayer, J.S., Bittner, R.A., Nikolić, D., Bledowski, C., Goebel, R., Linden, D.E.J., 2007. Common neural substrates for visual working memory and attention. NeuroImage 36, 441–453. https://doi.org/10.1016/j.neuroimage.2007.03.007

Mitchell, T.M., Shinkareva, S.V., Carlson, A., Chang, K.-M., Malave, V.L., Mason, R.A., Just, M.A., 2008. Predicting Human Brain Activity Associated with the Meanings of Nouns. Science 320, 1191–1195. https://doi.org/10.1126/science.1152876

Mottolese, C., Richard, N., Harquel, S., Szathmari, A., Sirigu, A., Desmurget, M., 2013. Mapping motor representations in the human cerebellum. Brain 136, 330–342. https://doi.org/10.1093/brain/aws186

Nishimoto, S., Vu, A.T., Naselaris, T., Benjamini, Y., Yu, B., Gallant, J.L., 2011. Reconstructing visual experiences from brain activity evoked by natural movies. Curr. Biol. 21, 1641–1646. https://doi.org/10.1016/j.cub.2011.08.031

Ortega, A., Frossard, P., Kovačević, J., Moura, J.M.F., Vandergheynst, P., 2018. Graph Signal Processing: Overview, Challenges and Applications. Proc IEEE 106, 808–828.

Poldrack, R.A., 2011. Inferring Mental States from Neuroimaging Data: From Reverse Inference to Large-Scale Decoding. Neuron 72, 692–697. https://doi.org/10.1016/j.neuron.2011.11.001

Poldrack, R.A., Halchenko, Y., Hanson, S.J., 2009. Decoding the Large-Scale Structure of Brain Function by Classifying Mental States Across Individuals. Psychol. Sci. 20, 1364–1372. https://doi.org/10.1111/j.1467-9280.2009.02460.x

Poldrack, R.A., Mumford, J.A., Schonberg, T., Kalar, D., Barman, B., Yarkoni, T., 2012. Discovering Relations Between Mind, Brain, and Mental Disorders Using Topic Mapping. PLoS Comput. Biol. 8, e1002707. https://doi.org/10.1371/journal.pcbi.1002707

Pulvermüller, F., Tomasello, R., Henningsen-Schomers, M.R., Wennekers, T., 2021. Biological constraints on neural network models of cognitive function. Nat. Rev. Neurosci. 22, 488–502. https://doi.org/10.1038/s41583-021-00473-5

Quian Quiroga, R., 2019. Plugging in to Human Memory: Advantages, Challenges, and Insights from Human Single-Neuron Recordings. Cell 179, 1015–1032. https://doi.org/10.1016/j.cell.2019.10.016

Richiardi, J., Eryilmaz, H., Schwartz, S., Vuilleumier, P., Van De Ville, D., 2011. Decoding brain states from fMRI connectivity graphs. NeuroImage, Multivariate Decoding and Brain Reading 56, 616–626. https://doi.org/10.1016/j.neuroimage.2010.05.081

Rosen, B.Q., Halgren, E., 2021. A Whole-Cortex Probabilistic Diffusion Tractography Connectome. eNeuro 8. https://doi.org/10.1523/ENEURO.0416-20.2020

Rosenberg, M.D., Scheinost, D., Greene, A.S., Avery, E.W., Kwon, Y.H., Finn, E.S., Ramani, R., Qiu, M., Constable, R.T., Chun, M.M., 2020. Functional connectivity predicts changes in attention observed across minutes, days, and months. Proc. Natl. Acad. Sci. 117, 3797–3807. https://doi.org/10.1073/pnas.1912226117

Schaefer, A., Kong, R., Gordon, E.M., Laumann, T.O., Zuo, X.-N., Holmes, A.J., Eickhoff, S.B., Yeo, B.T.T., 2018. Local-Global Parcellation of the Human Cerebral Cortex from Intrinsic Functional Connectivity MRI. Cereb. Cortex N. Y. NY 28, 3095–3114. https://doi.org/10.1093/cercor/bhx179

Schon, K., Newmark, R.E., Ross, R.S., Stern, C.E., 2016. A Working Memory Buffer in Parahippocampal Regions: Evidence from a Load Effect during the Delay Period. Cereb. Cortex 26, 1965–1974. https://doi.org/10.1093/cercor/bhv013

Schrimpf, M., Kubilius, J., Hong, H., Majaj, N.J., Rajalingham, R., Issa, E.B., Kar, K., Bashivan, P., Prescott-Roy, J., Geiger, F., Schmidt, K., Yamins, D.L.K., DiCarlo, J.J., 2020. Brain-Score: Which Artificial Neural Network for Object Recognition is most Brain-Like? bioRxiv 407007. https://doi.org/10.1101/407007

Selvaraju, R.R., Cogswell, M., Das, A., Vedantam, R., Parikh, D., Batra, D., 2020. Grad-CAM: Visual Explanations from Deep Networks via Gradient-based Localization. Int. J. Comput. Vis. 128, 336–359. https://doi.org/10.1007/s11263-019-01228-7

Shirer, W.R., Ryali, S., Rykhlevskaia, E., Menon, V., Greicius, M.D., 2012. Decoding Subject-Driven Cognitive States with Whole-Brain Connectivity Patterns. Cereb. Cortex 22, 158–165. https://doi.org/10.1093/cercor/bhr099

Sporns, O., 2018. Graph theory methods: applications in brain networks. Dialogues Clin. Neurosci. 20, 111–121.

Sporns, O., 2011. The Non-Random Brain: Efficiency, Economy, and Complex Dynamics. Front. Comput. Neurosci. 5. https://doi.org/10.3389/fncom.2011.00005

Springenberg, J.T., Dosovitskiy, A., Brox, T., Riedmiller, M., 2014. Striving for Simplicity: The All Convolutional Net.

Suárez, L.E., Richards, B.A., Lajoie, G., Misic, B., 2021. Learning function from structure in neuromorphic networks. Nat. Mach. Intell. 3, 771–786. https://doi.org/10.1038/s42256-021-00376-1

Tavor, I., Jones, O.P., Mars, R.B., Smith, S.M., Behrens, T.E., Jbabdi, S., 2016. Task-free MRI predicts individual differences in brain activity during task performance. Science 352, 216–220. https://doi.org/10.1126/science.aad8127

Urchs, S., Armoza, J., Moreau, C., Benhajali, Y., St-Aubin, J., Orban, P., Bellec, P., 2019. MIST: A multi-resolution parcellation of functional brain networks. MNI Open Res. 1, 3. https://doi.org/10.12688/mniopenres.12767.2

Varoquaux, G., Baronnet, F., Kleinschmidt, A., Fillard, P., Thirion, B., 2010. Detection of Brain Functional-Connectivity Difference in Post-stroke Patients Using Group-Level Covariance Modeling, in: Jiang, T., Navab, N., Pluim, J.P.W., Viergever, M.A. (Eds.), Medical Image Computing and Computer-Assisted Intervention – MICCAI 2010, Lecture Notes in Computer Science. Springer, Berlin, Heidelberg, pp. 200–208. https://doi.org/10.1007/978-3-642-15705-9_25

Varoquaux, G., Schwartz, Y., Poldrack, R.A., Gauthier, B., Bzdok, D., Poline, J.-B., Thirion, B., 2018. Atlases of cognition with large-scale human brain mapping. PLOS Comput. Biol. 14, e1006565. https://doi.org/10.1371/journal.pcbi.1006565

Wang, X., Liang, X., Jiang, Z., Nguchu, B.A., Zhou, Y., Wang, Y., Wang, H., Li, Y., Zhu, Y., Wu, F., Gao, J., Qiu, B., 2020. Decoding and mapping task states of the human brain via deep learning. Hum. Brain Mapp. 41, 1505–1519. https://doi.org/10.1002/hbm.24891

Yamashita, M., Yoshihara, Y., Hashimoto, R., Yahata, N., Ichikawa, N., Sakai, Y., Yamada, T., Matsukawa, N., Okada, G., Tanaka, S.C., Kasai, K., Kato, N., Okamoto, Y., Seymour, B., Takahashi, H., Kawato, M., Imamizu, H., 2018. A prediction model of working memory across health and psychiatric disease using whole-brain functional connectivity. eLife 7, e38844. https://doi.org/10.7554/eLife.38844

Yeo, B.T.T., Krienen, F.M., Sepulcre, J., Sabuncu, M.R., Lashkari, D., Hollinshead, M., Roffman, J.L., Smoller, J.W., Zöllei, L., Polimeni, J.R., Fischl, B., Liu, H., Buckner, R.L., 2011. The organization of the human cerebral cortex estimated by intrinsic functional connectivity. J. Neurophysiol. 106, 1125–1165. https://doi.org/10.1152/jn.00338.2011

Zhang, Y., Tetrel, L., Thirion, B., Bellec, P., 2021. Functional annotation of human cognitive states using deep graph convolution. NeuroImage 231, 117847. https://doi.org/10.1016/j.neuroimage.2021.117847

Zielinski, B.A., Gennatas, E.D., Zhou, J., Seeley, W.W., 2010. Network-level structural covariance in the developing brain. Proc. Natl. Acad. Sci. 107, 18191–18196. https://doi.org/10.1073/pnas.1003109107

