## supplementary figures for "Deep learning models of cognitive processes constrained by human brain connectomes"

***** **Corresponding Authors:**

**This PDF file includes:**

Eight Supplementary Figures: **Figure 2-S1** to **Figure 9-S1**

### **Supplementary Figures**

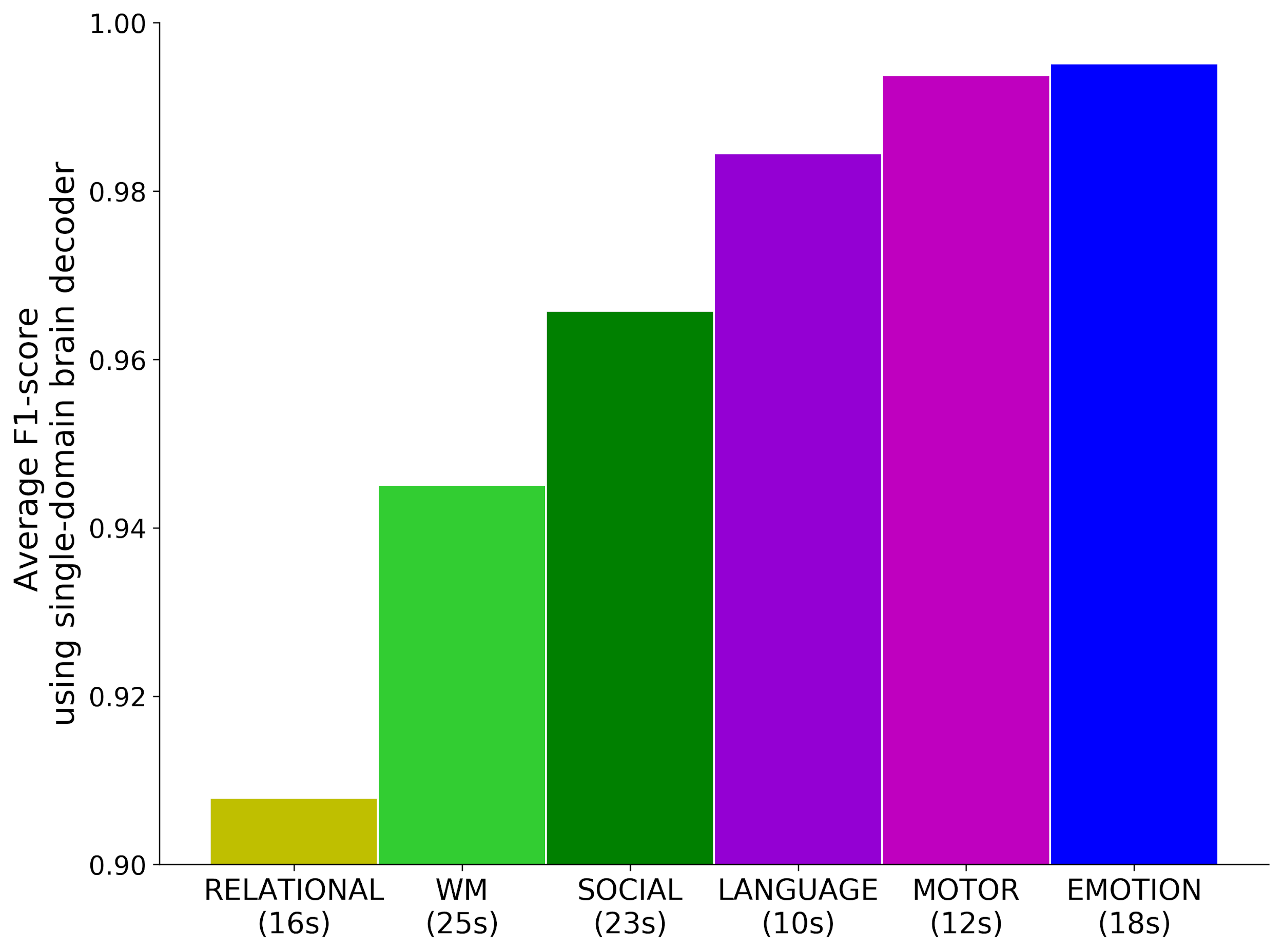

**Figure 2-S1. Decoding accuracy of single-domain brain decoders for each of the six cognitive domains.**

Variable temporal durations were used for the decoding models of each cognitive domain, according to the maximum length of event trials/blocks among task conditions, for instance 12s for MOTOR tasks and 25s for WM tasks. The same color scheme was used as Figure **2c**. Among the six cognitive domains, the emotion tasks (in blue, fearful face *vs* shape) and motor tasks (in magenta, distinguishing five types of body movements) were the most easily recognizable conditions, with F1-score reaching above 99%.

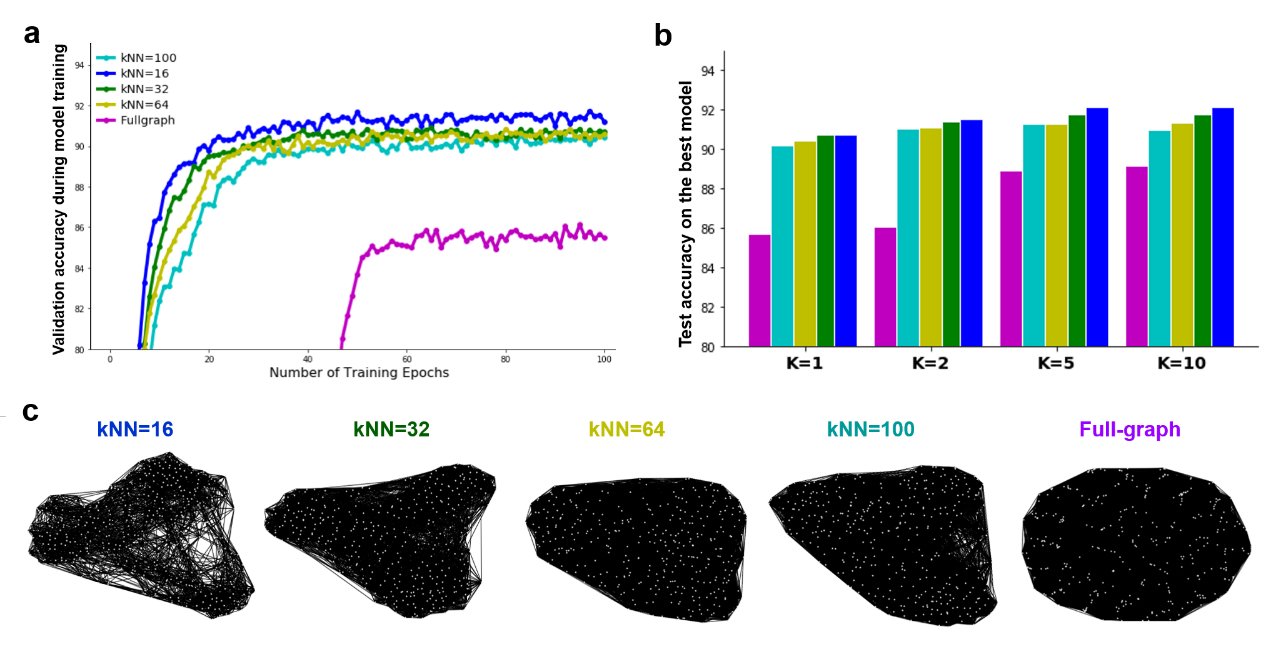

**Figure 2-S2. ChebNet decoding using densely connected functional graphs and graphs at different sparsity levels.**

The sparse brain graphs (in blue) significantly outperformed the original densely connected functional connectomes (in purple). During model training and evaluation, the edge-sparsified graphs showed much faster convergence speed and achieved much better decoding performance than the counterpart densely connected brain graphs. We trained separated decoding models at different K-orders and found that the highly sparsified graphs (e.g. kNN=16) achieved the best decoding performance at all K-orders. The decoding performance gradually decayed as the contention ratio increased (b). All edge-sparsified graphs were generated from the same group-wise connectivity matrix (i.e. full-graph in c) and then threshold at different sparsity levels by using k nearest neighbors of each node.

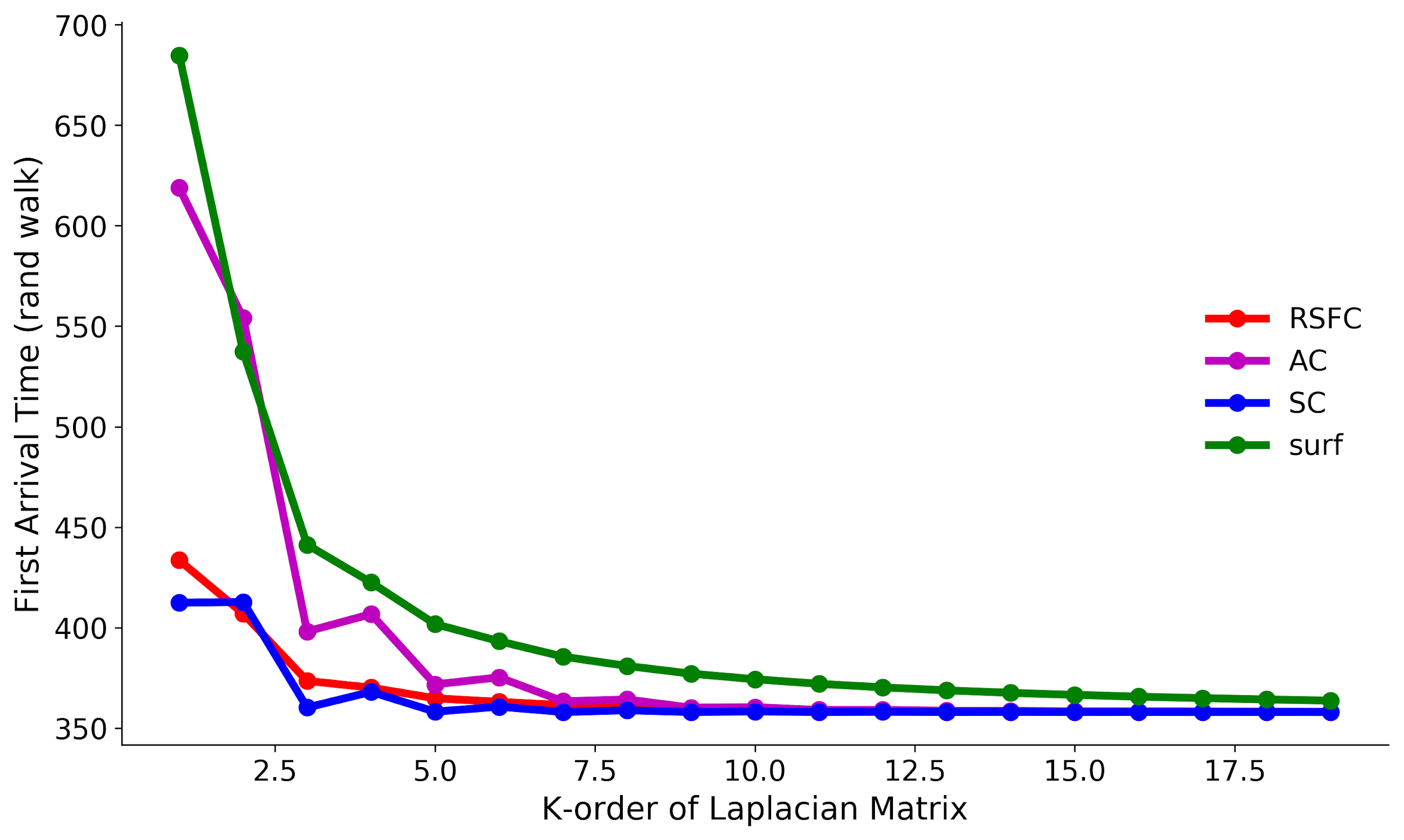

**Figure 3-S1. Information propagation speed as a function of the K-order.**

We used a metric in information theory, namely the first arrival time, to evaluate the information propagation speed on the brain graph, which represents the time required for the flow of the information first propagated from any source node to a given node. The expected value of first arrival time by taking random walks on the graph was calculated by, according to (Sakumoto et al., 2019):
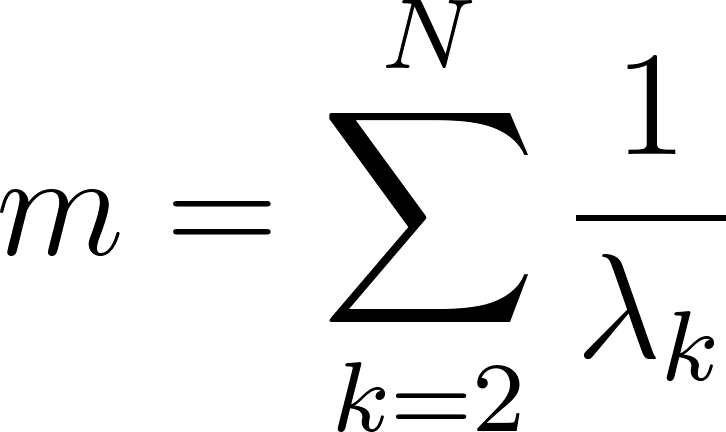
<https://www.codecogs.com/eqnedit.php?latex=m%3D%20%5Csum%5EN_%7Bk%3D2%7D%5Cfrac%7B1%7D%7B%5Clambda_%7Bk%7D%7D%20#0>[, where](https://www.codecogs.com/eqnedit.php?latex=m%3D%20%5Csum%5EK_%7Bk%3D2%2CN%7D1%2F%5Clambda_%7Bk%7D%20#0)
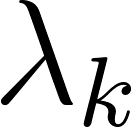
<https://www.codecogs.com/eqnedit.php?latex=%5Clambda_%7Bk%7D#0> [is the eigenvalues of the normalized Lapalican matrix.](https://www.codecogs.com/eqnedit.php?latex=m%3D%20%5Csum%5EK_%7Bk%3D2%2CN%7D1%2F%5Clambda_%7Bk%7D%20#0) As we increased the *K*-order, the first arrival time
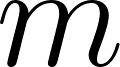
<https://www.codecogs.com/eqnedit.php?latex=m#0> was largely shortened, with the largest gradient appearing around $K=5$ and reaching the plateau around $K=10$ on all brain graphs. Four types of brain graphs were evaluated here, including spatial-graph (surf), structural-graph (SC), diffusion-graph (AC) and functional-graph (RSFC). Different sensitivity levels to the *K*-order were detected, such that information propagation on the spatial-graph was greatly impacted by the choice of *K*-order, followed by the diffusion-graph with a higher convergence speed. The functional-graph and structural-graph quickly converged to the plateau and followed a similar curve when increasing the *K*-order, except that higher randomized effect was observed on the structural graph at small *K*-orders.

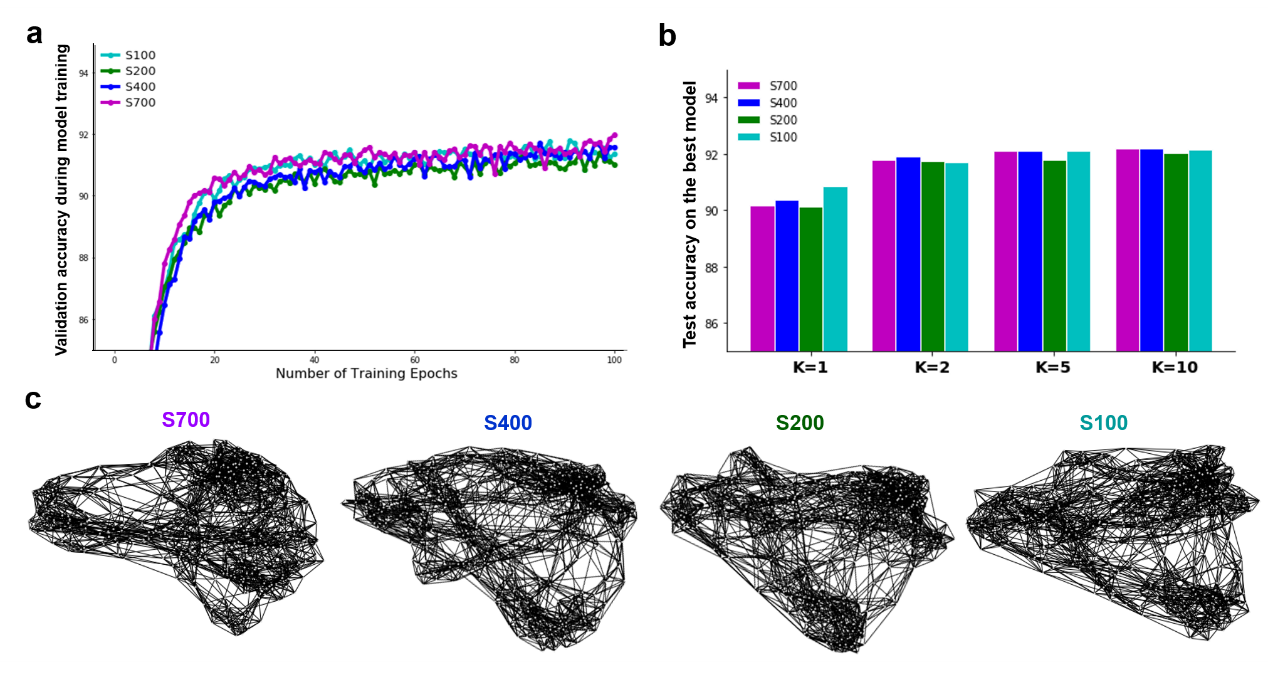

**Figure 3-S2. ChebNet decoding using brain graphs derived from different groups of training subjects.**

We constructed four different functional graphs by calculating the resting-state functional connectivity from 100, 200, 400 and 700 subjects in the training set. The decoding performance on the four brain graphs were evaluated at different K-orders. We found that the graph architecture derived from different groups of training subjects showed little impact on the cognitive decoding. Yet, for all brain graphs, the decoding performance gradually improved by increasing the K-order and plateaued at K=5.

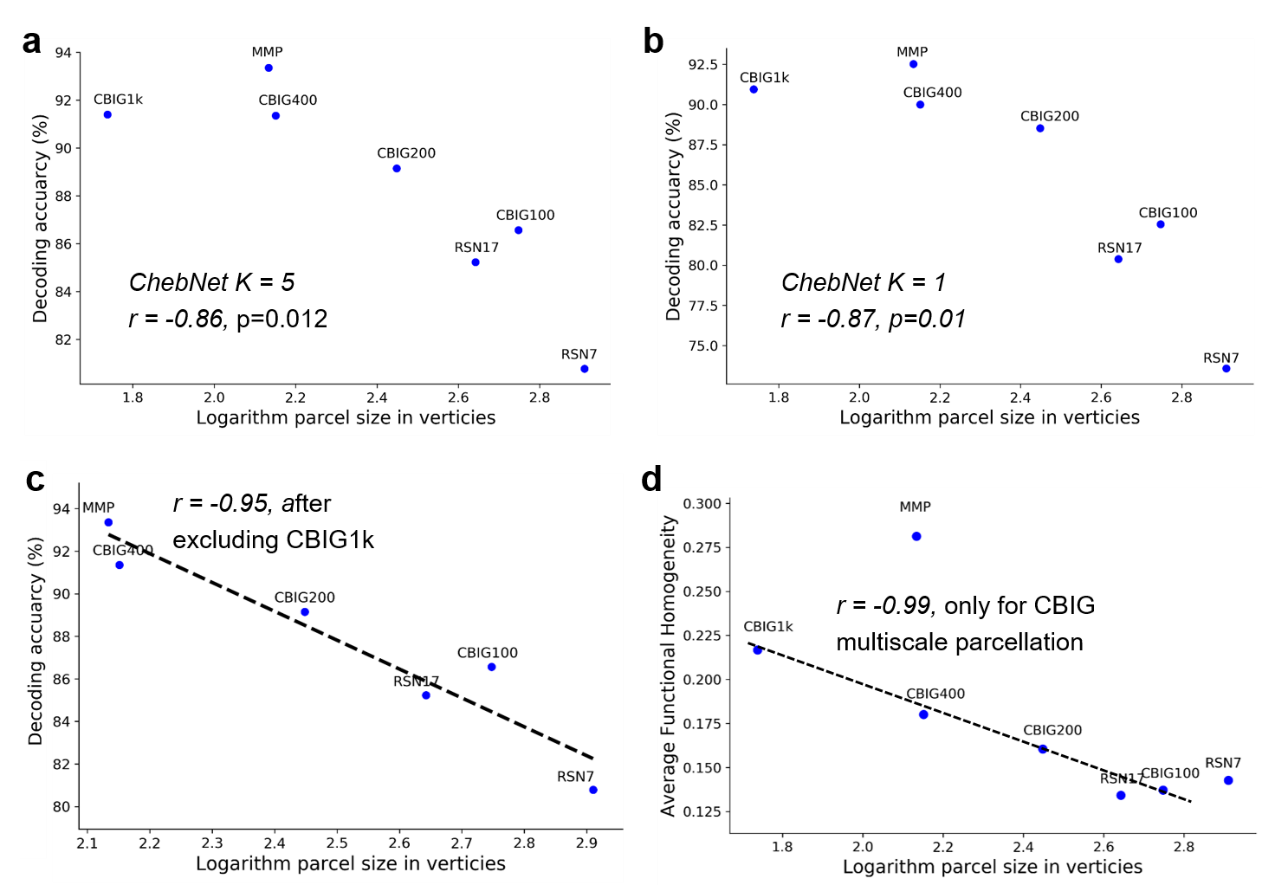

**Figure 5-S1. Decoding performance associated with parcel size across different brain atlases.**

We evaluated brain atlases at different resolutions for the construction of the brain graph and observed high variability in the decoding performance. The results showed that the parcel size had a significant impact on the decoding accuracy by using either ChebNet-K5 (**a**) or ChebNet-K1 (**b**) models. This effect became more significant when excluding the cases that the decoding model plateaued at more than 400 parcels (r =-0.95, p=0.003 in (**c**), excluding brain parcellation with 1000 regions). The parcel size also showed a significant impact on the functional homogeneity of each brain parcel such that finer-scale atlases (smaller parcel size) have higher functional homogeneity (**d**). For instance, among the Schaefer’s multiresolution brain parcellation (Schaefer et al., 2018), we found a significant association between parcel size and functional homogeneity (Pearson correlation r = -0.99, p=0.001 in d). Note that we did not include the Brainnetome atlas in this analysis because it is defined on volumetric data while the rest of atlases are defined on the cortical surface. The choice of volume space or cortical surface may influence the calculation of parcel size and functional homogeneity. Our results indicate that high-resolution brain atlases result in high functional homogeneity of brain parcels, which consequently achieve better performance for the decoding of cognitive states.

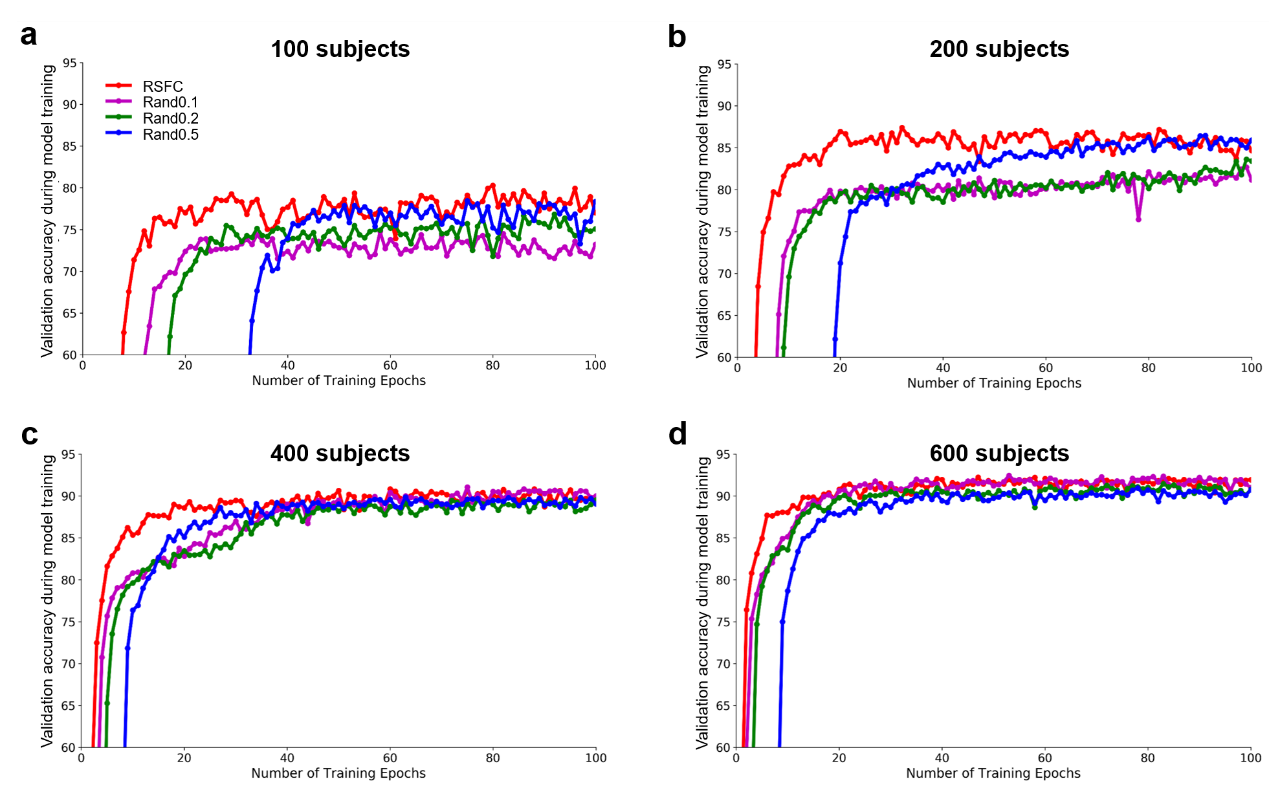

**Figure 7-S1. Brain decoding using ChebNet-K5 on randomized functional graphs and small datasets.**

We generated three sets of randomized brain graphs by randomly swapping a portion of edges in the functional graph while keeping the node degree unchanged. The network structures of the brain graphs were shown in Figure **6c**. Here, we found that the randomness of brain graphs showed a significant impact on the convergence speed during model training. When fixing the *K-*order, for example using the ChebNet-*K*5 model, and training the model on different sample sizes, for instance 100, 200, 400 and 600 subjects, the original functional graph (RSFC, in red) showed the fastest convergence during the training process at all cases, followed by the graphs of low (Rand0.1, in purple) and moderate randomization (Rand0.2, in green) and finally the highly randomized graph (Rand0.5, in blue). Interestingly, when only a small number of subjects were available, the highly randomized graph (Rand0.5) quickly caught up with the original functional graph during model training and eventually exceeded the decoding performance on lowly randomized graphs (e.g. Rand0.1). This effect gradually disappeared when using a sufficiently large sample size (i.e. more than 600 subjects).

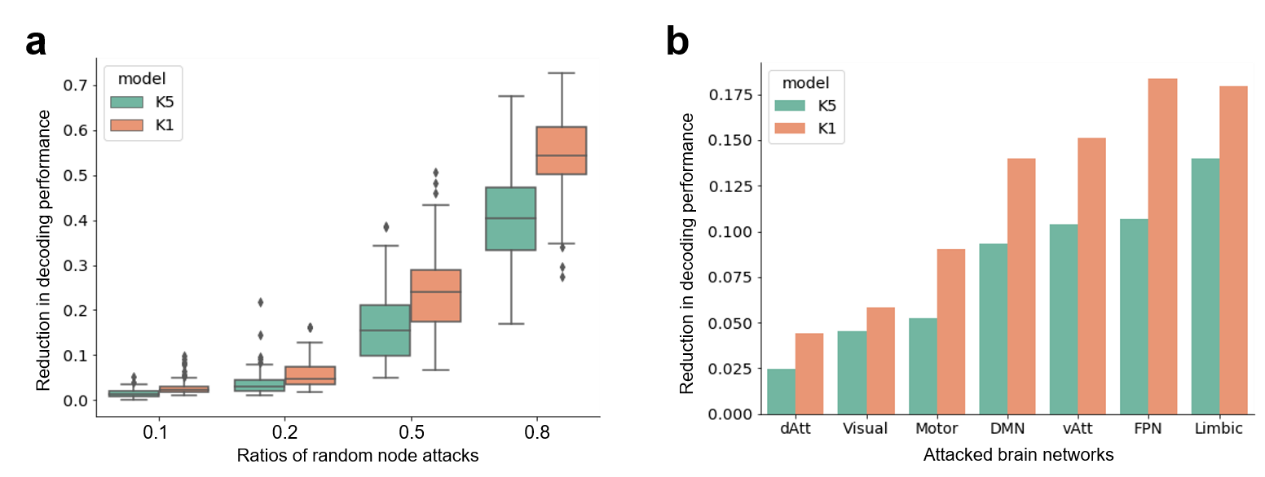

**Figure 8-S1. Decays in the decoding of WM tasks using ChebNet-*K*5 and ChebNet-*K*1 models.**

Compared to ChebNet-*K*5, the low-order model (e.g. *K=*1) was more vulnerable to random attacks on both brain regions (**a**) and networks (**b**). We simulated different types of brain lesions by silencing brain responses from randomly chosen nodes, ranging from 10% to 80% of brain parcels, or targeted regions from a specific brain network. We found that the decays of cognitive decoding were more severe in the ChebNet-*K*1 model at all levels for random node attacks (T-value=6.27, 4.64, 6.79 and 9.86 respectively, p-values<0.00001) and network attacks. The most affected areas consist of regions in the ventral visual stream and frontoparietal network and the most vulnerable networks are those related to cognitive control and attention, consistent with the findings on the ChebNet-*K*5 model.

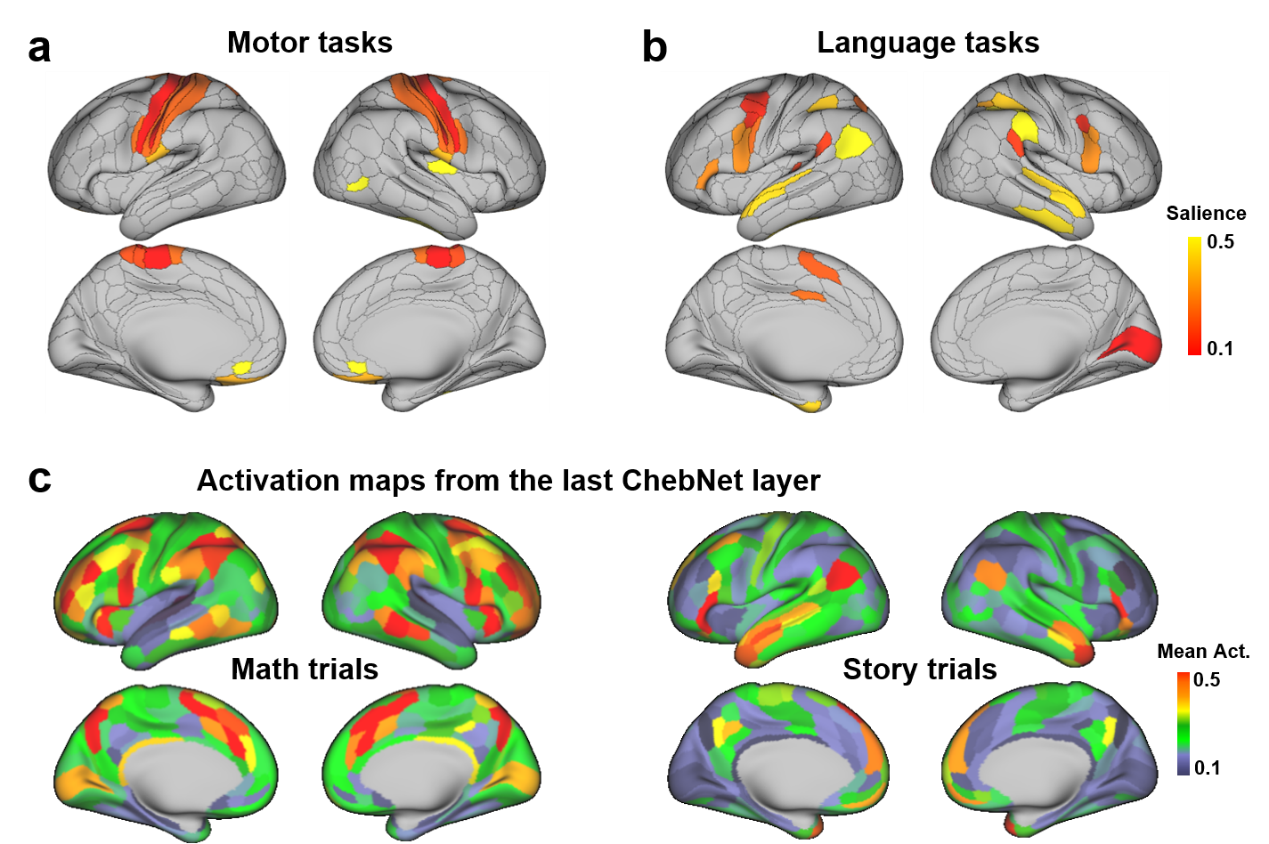

**Figure 9-S1. Saliency maps and activation patterns for the decoding of Motor and Language tasks.**

The model detected biologically meaningful and category-specific salient features on the Motor and Language tasks. For the Motor tasks (a), the model detected high salience in the primary motor and somatosensory cortices. For the Language tasks (b), the model identified salient regions in the primary auditory cortex and perisylvian language-related brain regions, consisting of inferior frontal gyrus (IFG), supramarginal gyrus/angular gyrus, and superior temporal gyrus (STG). Moreover, we mapped the graph representations in the last ChebNet layer onto the human brain for the mathematics and story trials and demonstrated distinct patterns of neural activity in the perisylvian language-related brain regions (c).

### **References**

Sakumoto, Y., Kameyama, T., Takano, C., Aida, M., 2019. Information Propagation Analysis of Social Network Using the Universality of Random Matrix. IEICE Trans. Commun. E102.B, 391–399. https://doi.org/10.1587/transcom.2018EBP3098

Schaefer, A., Kong, R., Gordon, E.M., Laumann, T.O., Zuo, X.-N., Holmes, A.J., Eickhoff, S.B., Yeo, B.T.T., 2018. Local-Global Parcellation of the Human Cerebral Cortex from Intrinsic Functional Connectivity MRI. Cereb. Cortex N. Y. NY 28, 3095–3114. https://doi.org/10.1093/cercor/bhx179

Zhang, Y., Tetrel, L., Thirion, B., Bellec, P., 2021. Functional annotation of human cognitive states using deep graph convolution. NeuroImage 231, 117847. https://doi.org/10.1016/j.neuroimage.2021.117847
